# Mechanistic insights into the evolution of DUF26-containing proteins in land plants

**DOI:** 10.1101/493502

**Authors:** Aleksia Vaattovaara, Benjamin Brandt, Sitaram Rajaraman, Omid Safronov, Andres Veidenberg, Markéta Luklová, Jaakko Kangasjärvi, Ari Löytynoja, Michael Hothorn, Jarkko Salojärvi, Michael Wrzaczek

## Abstract

Large protein families are a prominent feature of plant genomes and their size variation is a key element for adaptation in plants. Here we infer the evolutionary history of a representative protein family, the DOMAIN OF UNKNOWN FUNCTION (DUF) 26-containing proteins. The DUF26 first appeared in secreted proteins. Domain duplications and rearrangements led to the emergence of CYSTEINE-RICH RECEPTOR-LIKE PROTEIN KINASES (CRKs) and PLASMODESMATA-LOCALIZED PROTEINS (PDLPs). While the DUF26 itself is specific to land plants, structural analyses of Arabidopsis PDLP5 and PDLP8 ectodomains revealed strong similarity to fungal lectins. Therefore, we propose that DUF26-containing proteins constitute a novel group of plant carbohydrate-binding proteins. Following their appearance, CRKs expanded both through tandem duplications and preferential retention of duplicates in whole genome duplication events, whereas PDLPs evolved according to the dosage balance hypothesis. Based on our findings, we suggest that the main mechanism of expansion in new gene families is small-scale duplication, whereas genome fractionation and genetic drift after whole genome multiplications drive families towards dosage balance.

---

Gene duplication and loss events constitute the main factor of gene family evolution^1^. Duplications occur by two major processes, whole genome multiplications (WGM) and small-scale duplications (SSD), including tandem, segmental, and transposon-mediated duplications^2^. There appears to be two distinct modes of expansion, since the gene families that evolve through WGMs rarely experience SSD events^3^. The division is visible also on the functional level, since genes duplicated in WGMs are enriched for transcriptional and developmental regulation as well as signal transduction functions, whereas SSDs occur preferentially on secondary metabolism and environmental response genes^3^. The prevailing explanation for the phenomenon is dosage balance; in complex regulatory networks and protein complexes the stoichiometric balance between the different components needs to be preserved, and therefore selection acts against losses after WGMs and against duplications in SSDs^4^. In terms of sizes, the families retained after WGMs are stable across different species whereas highly variable families evolve through SSDs^5^, suggesting high turnover rates. However, these results have been obtained by analyzing the two extremes, the top families displaying pure WGM retention or SSD characteristics^3^ while most of the gene families likely evolve in an intermediate manner.

Plants and other eukaryotes have developed a wide range of signal transduction mechanisms for controlling cellular functions and to coordinate responses on cell, tissue, organ and organismal level. Plants in particular encode large gene families of secreted proteins^6–8^ and proteins with extracellular domains to respond to environmental and developmental cues but in most cases their functions are not known^4^. Signaling proteins with extracellular domains include receptor-like protein kinases (RLKs)^9,10^ and receptor-like proteins (RLPs)^11^. In RLKs, extracellular domains are involved in signal perception and protein-protein interactions^12^ while the intracellular kinase domain transduces signals to intracellular substrate proteins. The RLKs are involved in essential mechanisms including stress responses, hormone signaling, cell wall monitoring and plant development^12^. The large number of secreted proteins, RLKs and RLPs in plants may reflect their sessile lifestyle and need for meticulous monitoring of signals from other cells, tissues, or the environment. However, the large numbers make it difficult to dissect their conserved or specialized functions, and therefore a detailed understanding of their evolution in different plant lineages is needed. Phylogenetic relations between different groups of RLKs and RLPs have been described^9,13–16^ but only few have been physiologically and biochemically characterized^17^.

Here we carry out an in-depth analysis of one protein family involved in signaling to explore the dynamics and effect of the different duplication mechanisms on overall gene family evolution: the Domain of Unknown Function 26 (DUF26; Gnk2 or stress-antifungal domain)-containing proteins^18,19^. The DUF26 is an extracellular domain harboring a conserved cysteine motif (C-8X-C-2X-C) in its core. It is present in three types of plant proteins. The first class is CYSTEINE-RICH RECEPTOR-LIKE SECRETED PROTEINs (CRRSPs). CRRSPs form large subgroups in *Arabidopsis thaliana* and rice (*Oryza sativa*) but in most plants the size of the family has not been quantified. The best characterized CRRSP is Gnk2, a protein from *Gingko biloba* with single DUF26 which exhibits antifungal activity and acts as mannose-binding lectin *in vitro*^18,19^. Two maize CRRSPs have been shown to also bind mannose and participate in defence against a fungal pathogen^20^. The second class, CYSTEINE-RICH RECEPTOR-LIKE PROTEIN KINASES (CRKs), has a typical configuration of two DUF26 in the extracellular region and forms a large subgroup of RLKs in plants with 44 members encoded in the *Arabidopsis thaliana* genome. CRKs participate in the control of stress responses and development in Arabidopsis and in rice^21–31^. The third class of DUF26 domain-containing proteins is the PLASMODESMATA-LOCALIZED PROTEINS (PDLPs). PDLPs contain two DUF26 domains in their extracellular region and a transmembrane helix, but lack a kinase domain. They associate with plasmodesmata and regulate symplastic intercellular signaling^32^, are involved in pathogen responses^33^, systemic signaling^34^, control of callose deposition^35^ and are targets for viral movement proteins^36^. However, the precise biochemical functions of DUF26-containing proteins in plants remain unclear.

Tandem expansions are a driving force for diversification processes for example for F-Box proteins^37^, transcription factors^38^, as well as RLKs^16^ and RLPs^11^. These diversification processes include sub-functionalization, where paralogs retain a subset of their original ancestral functions, and neo-functionalization, where a protein acquires novel functions after duplication^38^. CRKs and CRRSPs typically exist in clusters on plant chromosomes^24^, suggesting relatively recent tandem expansions. This makes the DUF26-containing proteins perfect dataset for testing the power of sequence-based evolutionary investigations. We propose that CRKs and CRRSPs experienced both ancestral and recent, lineage-specific tandem duplications in different angiosperm lineages. In contrast to the general pattern of gene families expanding by small-scale duplication events, these gene families experienced significant expansion also during of after ancient whole genome duplication events. We combine phylogenetic analyses with experimental structural biology to gain insight into the evolution of DUF26-containing proteins in plants. While sequence analysis indicates that the DUF26 domain is specific to land plants, the domain shows strong structural similarity to fungal carbohydrate-binding lectins. Our structural analyses suggest that DUF26-containing proteins constitute a novel group of carbohydrate-binding proteins in plants. Consequently, sequence similarity alone is not sufficient evidence of orthologs, and lineage-specific protein family expansions can make translation of functional data between species difficult. Our results illustrate that a detailed understanding of the evolution of large protein families is a prerequisite for translating findings from model plants to different species and for dissecting conserved or specialized functions of protein family members.

## Results

### Identification and annotation of DUF26 genes

We selected 32 plant species representing major lineages of the plant kingdom for which high-quality genome assemblies are available and retrieved 1656 DUF26-containing gene models (Figure 1a, Table S1). Manual curation identified 322 gene models that required correction, demonstrating the necessity of manual validation of datasets for analysis of gene families (Figure S1). To further reduce the possible biases in annotation quality, we searched and identified 268 gene models *de novo* from genomic sequences (see Materials and Methods). Partial gene models and pseudogenes were excluded resulting in 1409 high-quality models included in subsequent analyses.

**Figure 1.**
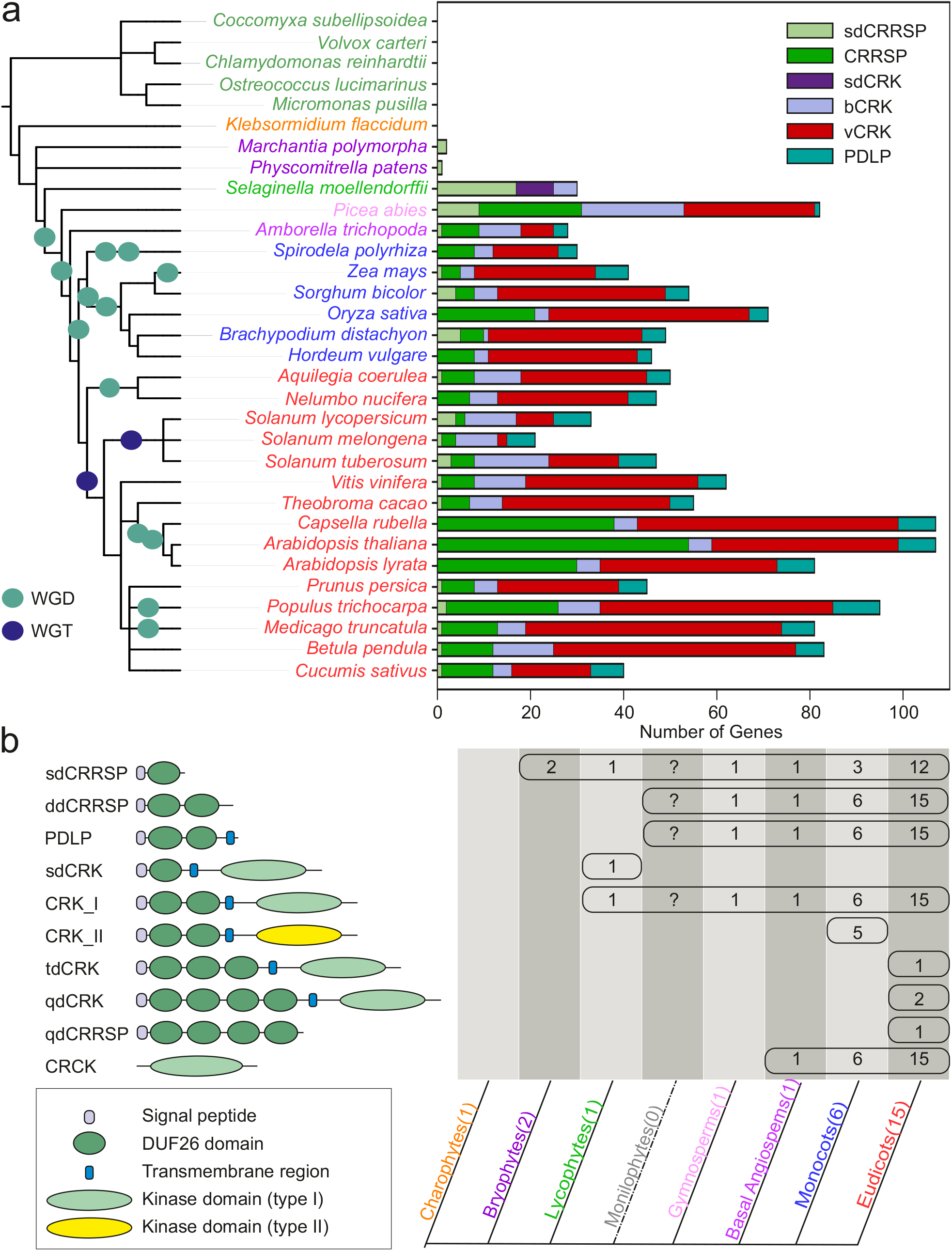
Overview and distribution of DUF26-containing genes in plants. **a)** DUF26-containing genes are absent from algae and charophytes but present in land plants. *Marchantia polymorpha* and *Physcomitrella patens* genomes encode sdCRRSPs. *Selaginella moellendorffii* possesses sdCRRSPs, sdCRKs and canonical CRKs. Seed plant (gymnosperm and angiosperm) genomes encode the whole set of DUF26-containing genes. CRKs were defined as basal group CRKs (bCRKs) or variable group CRKs (vCRKs) based on their phylogenetic positions. Whole genome duplication (WGD) events are presented with green circle and whole genome triplication (WGT) events with dark blue circle. Ferns were omitted from analyses due to lack of available genome assemblies. **b)** Overview of different domain compositions of proteins containing DUF26 in different plant lineages. The number of representative species in the analyses is given in brackets after the name of the group. Numbers in the table present the number of species in each lineage in which the domain structure was found. In abbreviations sd (single domain), dd (double domain), td (triple domain) and qd (quadruple domain) refers to the number of the DUF26 domains.

According to the PFAM protein domain database^39^, DUF26 is specific to embryophytes. We confirmed this by querying the genomes of the diatom *Phaeodactylum tricornutum*, five algae species, the charophyte *Klebsormidium flaccidum*, as well as fungi, insects and vertebrates (see Materials and Methods) and identified no DUF26 or DUF26-like domain among the species (Figure 1a).

### DUF26-containing proteins have diverse domain compositions

DUF26-containing proteins are grouped to three categories: CRRSPs, PDLPs and CRKs (Figure 1b). CRRSPs consist of a signal peptide (SP) followed by one or more DUF26 domains, separated by a short variable region. CRRSPs with a single DUF26 (sdCRRSPs) were identified from most land plants, including the early-diverging liverwort (*Marchantia polymorpha*) and moss (*Physcomitrella patens*) lineages (Figure 1). CRRSPs with two DUF26 domains (ddCRRSPs) were identified from vascular plants including the early-diverging lycophyte *Selaginella moellendorffii*; they represent the predominant type in all vascular plant genomes (Figure 1). Rice as well as *Brassicaceae* display lineage-specific evolution with a large number of ddCRRSPs while sdCRRSPs are absent (Figure 1a and S2).

CRKs contain a SP, two DUF26 domains, and a transmembrane region (TMR) followed by an intracellular protein kinase domain. Similar to ddCRRSPs, CRKs were identified from vascular plants but not from bryophytes (Figure 1a). The CRKs likely emerged as the result of a fusion of sdCRRSPs with TMR and kinase domain from LRR_clade_3 RLKs in the common ancestor of vascular plants^15^, since the *Selaginella* genome uniquely encodes single DUF26 CRKs (sdCRKs; Figure 1b). The two-domain configuration is stable, since only few CRKs from eudicot plants contain more than two DUF26 domains.

Finally, PDLPs are composed of a SP, two DUF26, and a transmembrane region (TMR) followed by a 10-15 amino acid (AA)-long cytoplasmic extension and they were identified from all seed plants. Within the angiosperms, we also identified several CRKs lacking SP, extracellular region and transmembrane domain. These are subsequently referred to as CYSTEINE-RICH RECEPTOR-LIKE CYTOPLASMIC KINASEs (CRCKs).

### Evolution of CRKs, PDLPs and ddCRRSPs from small sdCRRSPs

To investigate the relationships between CRRSPs, CRKs and PDLPs, we estimated phylogenetic trees using full length AA sequences translated from gene models with intact DUF26 domains (Figure 2a and b). As a result of the different domain compositions only the DUF26-containing region aligned across all sequences. Due to their high sequence divergence CRCKs, DUF26-containing gene models from bryophytes and monocot CRKs with a different intracellular protein kinase domain were excluded from the alignment (see below). Overall, a phylogenetic tree for DUF26-containing proteins based on a filtered amino acid sequence alignment split into two distinct groups, a basal group α and a variable group β (Figures 2a and b), where α is paraphyletic with respect to β. In order to increase the number of informative sites and thus obtain better resolution, we estimated separate phylogenetic trees for both groups (See Methods; Figures S2a and b); the subgrouping observed within the basal α- and variable β-groups was present there as well as in the trees estimated for each sub-family of DUF26-containing proteins (Figures S2c-e). To study gene family evolution we reconciled the gene trees with the species tree, and estimated ancestral gene contents and duplication and loss events for the sub-families in eleven species (see Materials and Methods; Figure S3). To identify significant expansions we fitted birth-death rate models for DUF26-containing protein families and compared the rates against different computationally derived gene families (orthogroups) for RLKs, all protein kinases, and plasmodesmal proteins^40^ using Badirate^41^ (see Materials and Methods). Finally, we assessed selective pressure by estimating amino acid conservation patterns around the main cysteine-motif of the DUF26 domains for major subclades within the α- and β-groups (Figures 2c). While most conserved positions within DUF26-A and -B are either conserved in all DUF26-containing proteins or specific to individual types, we were able to identify conserved sites specific to the α- or β-clades (Figures 2c).

**Figure 2.**
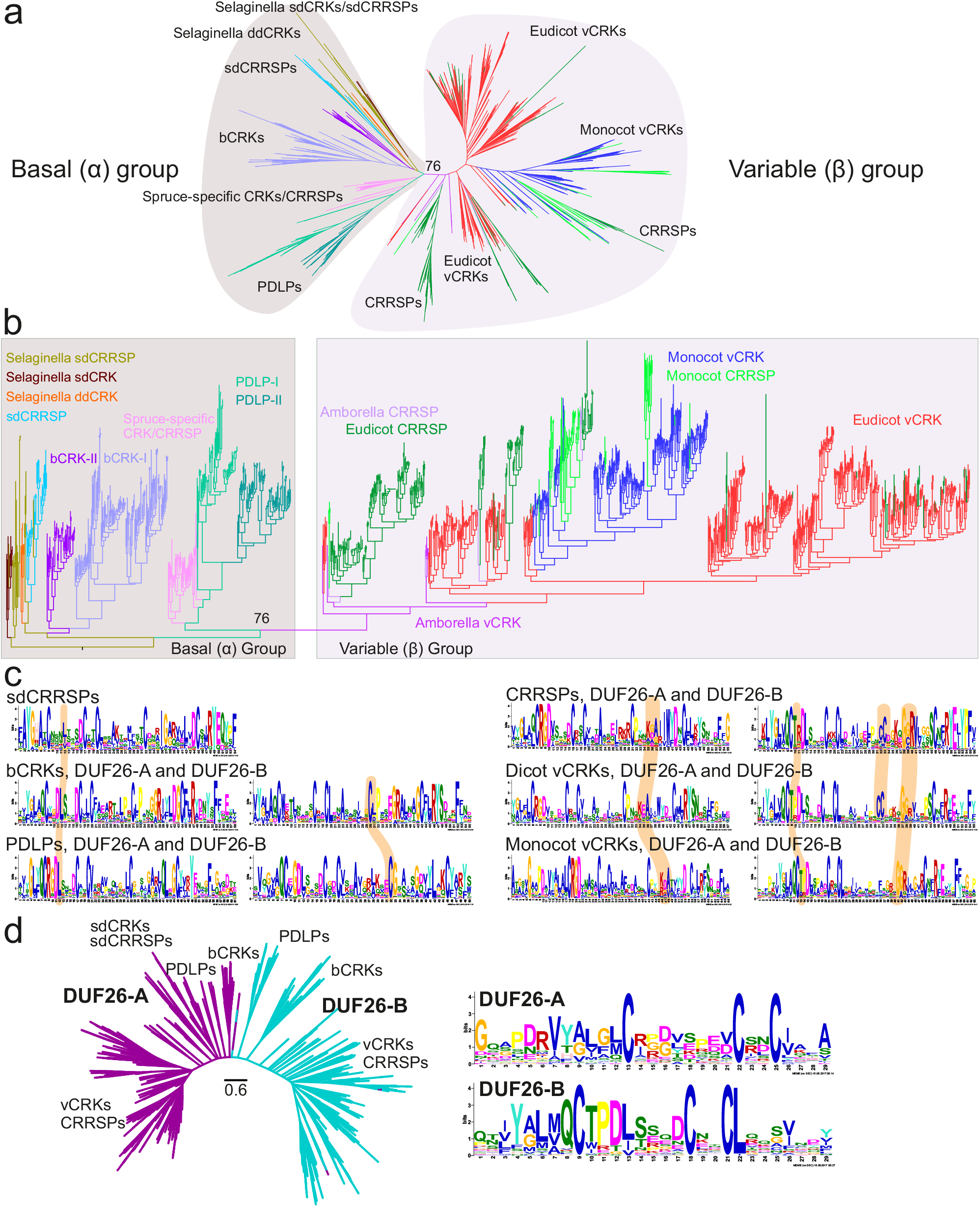
Phylogenetic tree of CRRSPs, CRKs and PDLPs. **a)** The phylogenetic tree was estimated with the maximum-likelihood method using all high quality full-length DUF26-containing sequences from lycophytes onwards. CRCKs and concA-CRKs were excluded while GNK2 from *Gingko biloba* was included. Overall, DUF26-containing genes split into basal and variable group. Detailed phylogenetic trees with bootstrap support (1000 replicates) and filtered sequence alignments are available at http://was.bi?id=IaroPa (full tree), http://was.bi?id=wpEHGt (basal group separately) and http://was.bi?id=aIJe_D (variable group separately). **b)** The same phylogenetic tree as in panel a rooted to ancestral sdCRRSPs and sdCRKs from *Selaginella moellendorffii* showing that the variable group branches out from the basal group. **c)** The MEME figures present the conservation pattern of amino acid positions around the main cysteine motif within the DUF26 domains for sdCRRSPs, bCRKs and PDLPs from the basal group and CRRSPs and vCRKs from the variable group. The features specific only to genes either in the basal group or in the variable group are highlighted. **d)** The DUF26-A and DUF26-B domains are clearly separated in an unrooted phylogenetic tree containing DUF26 domain sequences. The MEME figures present differences in the conservation of the AA sequence surrounding the conserved cysteines in DUF26-A and DUF26-B.

The α-group is likely older, containing sequences from all vascular plants. Proteins in this group are conserved in sequence level and identification of putative orthologs from different species is frequently possible. Purifying selection, i.e. stabilizing selection by selective removal of (deleterious) variations, is likely the main force acting on this clade, as suggested by low *d_N_*/*d_S_* values (one-rate model for whole groups: bCRK-I 0.184, bCRK-II 0.192, PDLPs 0.267, sdCRRSPs 0,162, CRCK 0.134; more flexible model with branch-specific *d_N_*/*d_S_* within each group yielded similar results). The subgroups within the basal α-group have evolved independently but their DUF26 domains share a number of features which distinguish them from the members of the variable β-group. These distinguishing features include a leucine or isoleucine residue in the fourth position after the first cysteine in the DUF26-A and the position of the fourth cysteine in the DUF26-B (Figure 2c). The sdCRRSPs appear to be the most ancient type of DUF26 genes in land plants, since the sdCRRSPs are located close to the root of the α-group (Figure 2b) and form a monophyletic subclade at the root of the CRRSP tree (Figure S2c). Furthermore, the sdCRRSPs are present in various early diverging plant lineages such as the gymnosperm *Ginkgo biloba* (including Gnk2, the best studied sdCRRSP^18,19^) and the liverwort, *Marchantia polymorpha* (Figure S2f). The turnover rates of sdCRRSPs do not differ from those of all gene families and show lineage-specific expansions in early diverging species (Figure S3a).

The placement of *Selaginella* sdCRKs to the root of the CRK phylogeny (Figure S2d) and as sister to sdCRRSPs in the α-group (Figure 2b) suggests an ancient origin. Our analyses indicate that the DUF26 domain likely has duplicated after fusing with TMR and kinase domain, thus establishing the typical double-DUF26 CRK configuration found in seed plants (Figure S2d). Following the duplication, the two DUF26 domains diverged into distinct forms, DUF26-A and DUF26-B, which are evolutionarily conserved (Figure 2d). Overall, CRKs have expanded significantly in the branches leading to lycophytes and to angiosperms compared to all RLKs (Figure 3a), and compared to all protein kinases they expanded significantly in the branch from the ancestral node of lycophyte *Selaginella* to angiosperms (Figure S4a). In all of these branches plants have experienced several WGDs^42^, suggesting that the ancestral CRKs have either been preferentially retained after WGMs„ or they have had a tandem birth rate that is higher than the death rate following WGMs.

**Figure 3.**
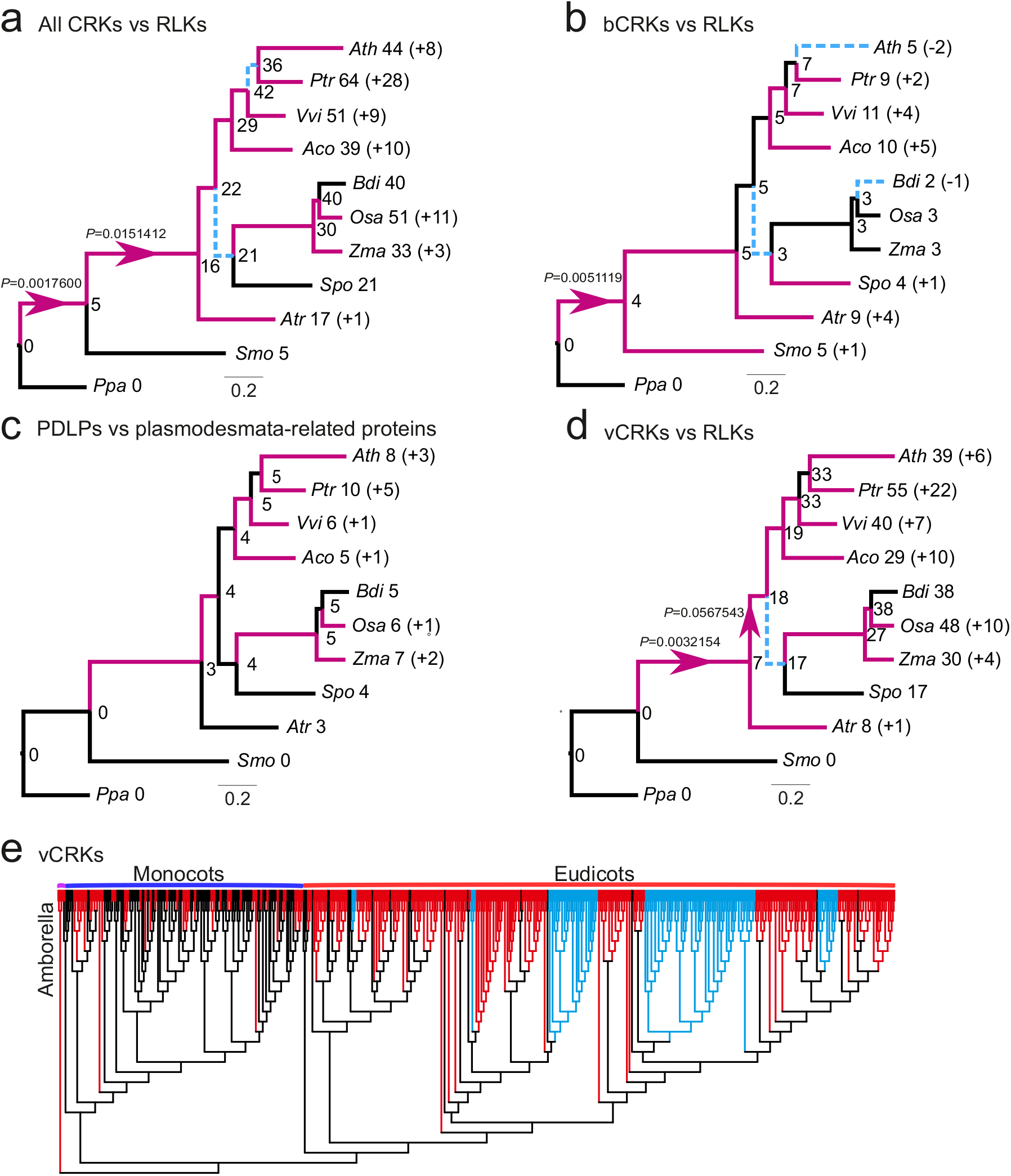
Comparison of evolutionary rates between gene families. Analyses were carried out with Badirate for eleven species (*Physcomitrella patens*, *Selaginella moellendorffii*, *Amborella trichopoda*, *Arabidopsis thaliana*, *Populus trichocarpa*, *Vitis vinifera*, *Aquilegia coerulea*, *Spirodela polyrhiza*, *Zea mays*, *Oryza sativa* and *Brachypodium distachyon*). Neutral branches are reported as bold black lines; branches involving gene family expansion are reported as bold purple lines and branches with contraction as blue dashed lines. Branches with a significant differences (false discovery rate adjusted p<0.05) to birth-death rate model estimates are marked with arrows. Node labels present the ancestral gene family sizes estimated by Badirate. Tip labels contain species abbreviations and the change in numbers compared to the most recent ancestral node. **a)** All CRKs compared to other receptor like kinases (RLKs). **b)** bCRKs compared to RLKs. **c)** PDLPs compared to other plasmodesmata related orthogroups. **d)** vCRKs compared to RLKs. **e)** Phylogenetic maximum-likelihood tree showing differences in lineage specific expansions in monocot and dicot vCRKs following the split of *Amborella trichopoda*. Species-specific expansions (at least two genes from same species) are marked with red and clades including sequences from only *Brassicaceae* or *Solanaceae* are marked with blue.

A monophyletic group of CRKs with representatives from gymnosperms and angiosperms is located near the base of the CRK phylogeny (Figure S2d) and belongs to the α-group (Figure 2a and b). This group likely represents the ancient CRKs in seed plants and will be subsequently referred to as basal CRKs (bCRKs). Following the initial innovation in ancestral vascular plants the group has evolved at rates similar to comparable orthogroups containing all protein kinases or all RLKs (Figure 3b. S3b and S4b). The bCRKs split into two distinct subgroups, bCRK-I and bCRK-II (Figure 2b and S5), both of which are present in gymnosperms and angiosperms, suggesting diverging duplicates in early seed plants. The larger bCRK-I subclade further divides into distinct branches with tandemly duplicated *Amborella* bCRKs at their roots (Figure S5, S6a-b) suggesting rapid differentiation after tandem duplication in ancestral angiosperms. The lineage-specific size of the bCRK-I branch is conserved, except for an expansion specific to *Solanaceae*. The small bCRK-II subclade is interestingly absent from *Brassicaceae*.

PDLPs are found in seed plants but not bryophytes or lycophytes. PDLPs belong to the α-group (Figure 2a and b) and represent the most conserved class of DUF26-containing genes. As such, they do not display different expansion rates compared to plasmodesmata-related orthogroups^40^ (Figure 3c). PDLPs split into two branches, PDLP-I and PDLP-II (Figure S2e), which both contain eudicot and monocot PDLPs, suggesting that the divergence occurred already in common ancestral angiosperms. The PDLP-II branch further divides into two angiosperm-specific branches with *Amborella trichopoda* sequences at their roots, whereas the PDLP-I branch can be traced back to a single *Amborella trichopoda* PDLP. PDLPs and ddCRRSPs originate from the loss of kinase domains and/or TMRs from CRKs. This two-step process is supported by an atypical PDLP from *Amborella trichopoda* which is located at the root of the main ddCRRSP clade (Figure S7). The timing of the event cannot be inferred, since it is unclear whether gymnosperms also contain members of the PDLP-I branch. However, a database search identified one partial gene model, a candidate PDLP from the fern *Marsilea quadrifolia*^43^ lacking a transmembrane region (see Materials and Methods). This putative fern PDLP shows high similarity to PDLPs and places to the root of PDLPs in a phylogenetic tree estimated from PDLPs and CRKs (Figure S8).

A group of spruce-specific CRKs (spruce vCRKs) belongs to the α-group (Figure 2a and b) and are more related to PDLPs than other CRKs. They form a distinct group between bCRKs and a large group of angiosperm CRKs, the “variable CRK clade” (vCRKs; Figure 2a and b, S2d, S3d). These angiosperm vCRKs form the β-group together with ddCRRSPs and atypical monocot sdCRRSPs (Figure 2a and b). These CRRSPs likely evolved from vCRKs through the loss of TMR and kinase domains and, in case of sdCRRSPs, also of the DUF26-B domains. The β-group is less conserved compared to the more ancient α-group and branches into two eudicot-specific groups and one monocot-specific group with a small group of *Amborella tiichopoda* vCRKs at the root of the clade. Still, there are some conserved positions surrounding the main cysteine motif that distinguish members of the β- from the α-group, for example a conserved threonine following the first cysteine in DUF26-B (Figure 2c). Unlike proteins in the α-group, CRRSPs and vCRKs in the β-group have undergone several independent tandem expansions in different plant taxa (Figure 3d, 3e, S3d, S4c, S6c) and expanded significantly during the diversification of monocots and dicots. CRRSPs in the β-group are not monophyletic, suggesting several independent birth events resulting from partial duplications of vCRKs. Hence, expansion rates and extrapolation of ancestral gene counts for ddCRRSPs could not be reliably predicted (Figure S3e). Lineage-specific expansions in the β-group will make identification of orthologs challenging.

### Plant DUF26 domains form conserved tandem assemblies and are structurally related to fungal lectins

The high sequence divergence of the DUF26 proteins in different plant lineages and the strong lineage-specific expansions raise the question whether their overall structure is conserved and what elements distinguish the more closely related members of this protein family. The consensus DUF26 (PF01657) domain as defined in PFAM comprises ~90-110 amino-acids and contains the conserved cysteine motif C-8X-C-2X-C. Structural information is currently available only for the sdCRRSP Gnk2^19^ but not for proteins with a double DUF26 configuration, such as ddCRRSPs, CRKs and PDLPs. Mechanistic constraints restrict the evolution of protein structures, and therefore understanding structural conservation can provide essential clues for protein function. Furthermore, selection patterns may differ between a young and lineage-specific gene and an evolutionarily conserved gene.

Thus, we defined the structural relationship of tandem DUF26 domains by determining crystal structures of the AtPDLP5 (residues 26-241) and AtPDLP8 (21-253) ectodomains to 1.25 and 1.95 Å resolution, respectively (Table S2). Individual DUF26 domains feature two small α-helices folding on top of a central anti-parallel β-sheet (Figure 4a). The PDLP5 DUF26-A domain is found to be N-glycosylated at positions Asn69 and Asn132 in our crystals (Figure 4a). The secondary structure elements of DUF26 are covalently linked by three disulfide bridges, formed by six conserved Cys residues, part of which belong to the C-8X-C-2X-C motif (Figures 2d, 4a). We have previously suggested that tandem DUF26-domain containing proteins could be involved in ROS or redox sensing^24,26^. To assess the functional roles of the invariant disulfide bridges in PDLPs, we mutated the partially solvent exposed PDLP5^Cys101^, PDLP5^Cys148^ and PDLP5^Cys191^ to alanine. While the wild-type PDLP5 ectodomain behaves as a monomer in solution (Figure S9), the mutant proteins tend to aggregate in our biochemical preparations (Figure S9) and display reduced structural stability in thermofluor assays (Figure S10, see Materials and Methods). These experiments and our crystallographic data (Figure 4a) together suggest that the conserved disulfide bonds in PDLPs and potentially in other DUF26-domain containing proteins are involved in structural stabilization rather than redox signaling.

**Figure 4:**
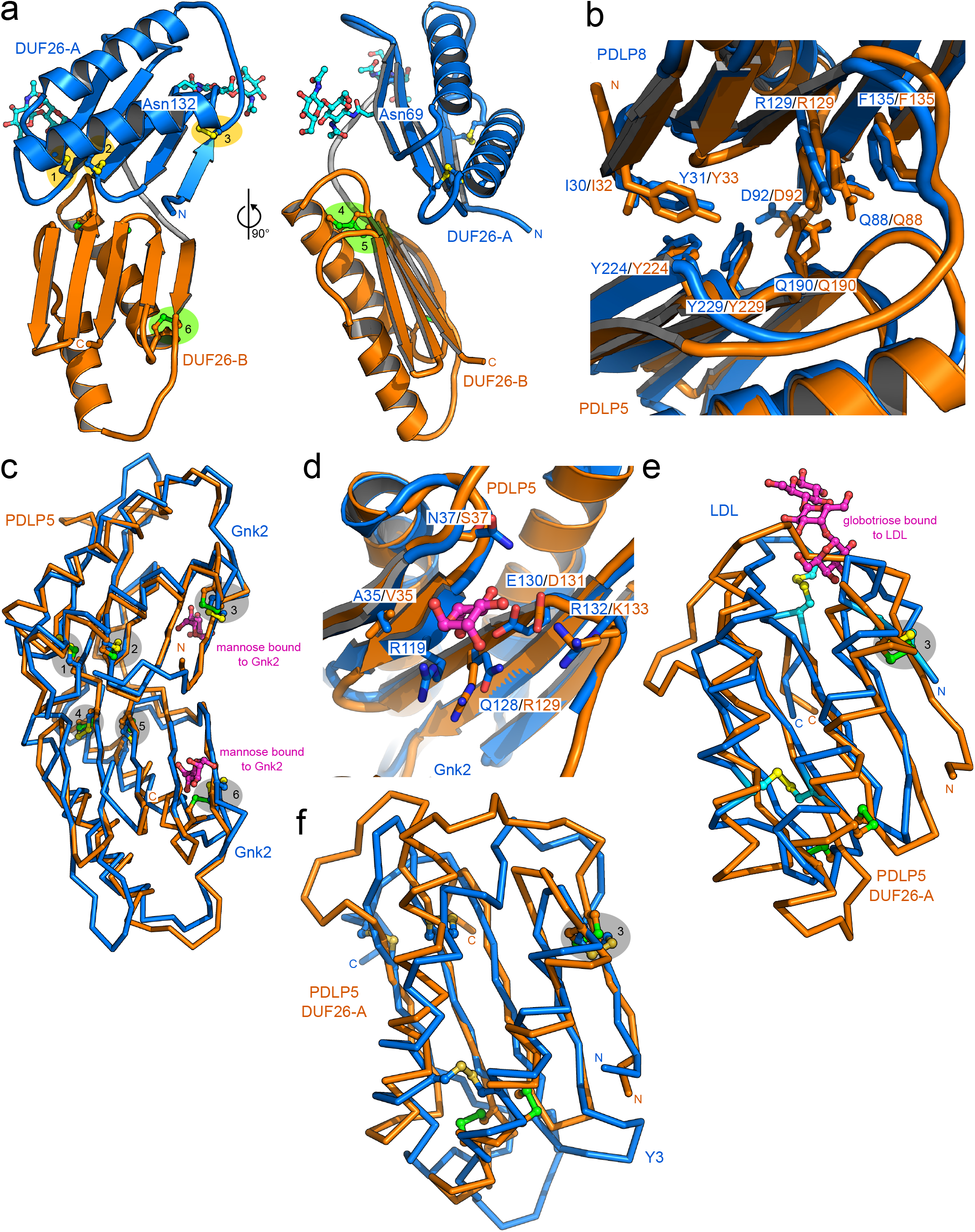
The crystals structures of the PDLP5 and PDLP8 ectodomains reveal a conserved tandem architecture of two lectin-like domains. **a)** Overview of the PDLP5 ectodomain. The two DUF26 domains are shown as ribbon diagrams, colored in blue (DUF26-A) and orange (DUF26-B), respectively. N-glycans are located at Asn69 and Asn132 of DUF26-A and are depicted in bonds representation (in cyan). The DUF26-A and DUF26-B domains each contain 3 disulfide bridges labeled 1 (Cys89-Cys98), 2 (Cys101-Cys126), 3 (Cys36-Cys113), 4 (Cys191-Cys200), 5 (Cys203-Cys228) and 6 (Cys148-Cys215). **b)** Close-up view of the DUF26-A – DUF26-B interface in PDLP5 (orange) and PDLP8 (blue), shown in bonds representation. **c)** Superimposition of the Gnk2 extracellular DUF26 domain (PDB-ID 4XRE) with either PDLP5 DUF-26A (r.m.s.d. is ~1.4 Å comparing 100 aligned C_α_ atoms) or PDLP5 DUF26-B (r.m.s.d. is ~2.0 Å comparing 93 corresponding C_α_ atoms). Corresponding disulfide bridges shown in bonds representation (PDLP5 in green, Gnk2 in yellow) are highlighted in grey. Gnk2-bound mannose is shown in magenta (in bonds representation). **d)** Close-up view of the residues involved in the binding of mannose of Gnk2 (bonds representation, in blue and magenta, respectively) and putative residues involved in substrate binding of PDLP5 DUF26-A (in orange). **e)** The fungal LDL DUF26 domain (C_α_ trace, in blue; PDB-ID 4NDV) and PDLP5 DUF26-A (in orange) superimposed with an r.m.s.d. of ~2.4 Å comparing 75 aligned C_α_ atoms). Disulfide bridges (LDL in yellow and PDLP5 in green; aligned disulfide bridges highlighted in grey) and the LDL bound globotriose (magenta) are shown in bonds representation. **f)** C_α_ traces of the structural superimposition of the fungal Y3 protein (PDB-ID 5V6I) and PDLP5 DUF26-A (r.m.s.d. is ~2.6 Å comparing 78 corresponding C_α_ atoms). Disulfide bridges of Y3 (yellow) and PDLP5 DUF26-A (green) are shown alongside, one corresponding disulfide pair is highlighted in gray.

The N-terminal DUF26-A (PDLP5 residues 30-132) and the C-terminal DUF26-B (residues 143-236) domains are connected by a structured loop (residues 133-142) and make extensive contacts with each other (Figure 4a). The resulting ectodomain has a claw-like shape with the β-sheets of DUF26 A and B facing each other (Figure 4a). The DUF26-A and B domains in PDLP5 and 8 closely align, with root mean square deviations (r.m.s.d.s) of 1.6 and 1.2 Å when comparing 89 corresponding C_α_ atoms, respectively (Figure S11a). Overall, DUF26-A is considerably more variable than DUF26-B on the sequence level (Figure 2d). The DUF26-A and B domains in PDLP5 and PDLP8 have 24 % and 30 % of their residues in common, most of which map to the hydrophobic core of the domain (including the six cysteine residues forming intramolecular disulfide bonds) and to the DUF26-A – DUF26-B interface (Figure 4b). This interface is formed by a line of aromatic and hydrophobic residues originating from the proximal face of the β-sheet in DUF26-A and B (Figure 4b, Supplementary Figure 12). Importantly, many of the interface residues are strongly conserved among different PDLPs, but also among CRKs and ddCRRSPs (Figure S12). Consistently, the ectodomains of PDLP5 and PDLP8 belonging to different phylogenetic clades (Figure S8) closely align with an r.m.s.d. of ~1.6 Å when comparing 198 corresponding C_α_ atoms (Figure S11b). Together, these observations suggest that evolutionarily distant DUF26 tandem proteins likely share the conserved three-dimensional structure.

The physiological ligands for PDLPs are currently unknown. For this means, we performed structural homology searches^44^ to obtain insights into the biochemical function of plant DUF26 domains (See Materials and Methods). The top hits include the single DUF26 domain protein ginkbilobin-2 (Gnk2) from *Ginko biloba*^19^. Despite their moderate sequence similarity, the overall fold of Gnk2 and PDLP5 DUF26-A and B as well as their disulfide-bond arrangement is fully conserved (Figure 4c). Notably, Glu130 and Arg132 implicated in mannose binding in Gnk2 are replaced by Asp131 and Lys133 in the DUF-A of PDLP5, respectively (Figure 4d). A similar pocket is found in the DUF-A domain of PDLP8, but not in the DUF-B domains of either PDLP5 or 8. Despite these structural homologies of Gnk2, PDLP5 DUF-A and PDLP8 DUF-A, we could not detect binding of mannose to the isolated PDLP5 ectodomain *in vitro* (Figure S13a). We also tested other water soluble cell wall derived carbohydrates, but were not able to detect any binding to the PDLP5 ectodomain (Figure S13b). The PDLP5 DUF26 domains share significant structural homology not only with the plant Gnk2, but also with two fungal lectins, the α-galactosyl-binding *Lyophyllum decastes* lectin (LDL)^45^ and a glycan-binding Y3 lectin from *Coprinus comatus*^46^. Both proteins closely align with the plant DUF26 domain, and share one of the three disulfide bridges (Figure 4e-f). The surface areas involved in globotriose and glycan binding, respectively, are not conserved in PDLPs, but the structural similarity of plant DUF26 domains with different eukaryotic lectins could suggest a common evolutionary origin and a potential role as carbohydrate recognition modules^45^.

We next explored potential binding sites in the two molecules by identifying regions under positive or purifying selection that could be indicative of domains involved in protein-protein interactions or ligand perception. Analysis of site-wise selection for orthologs of PDLP5 and PDLP8 in their structural context yielded low ω values, indicating strong conservation of residues buried inside the DUF26 domain fold, while more variable residues (under more relaxed selection) appear on the surface of the structure (Figure 5a). The high variability of the surface of the PDLP5 and PDLP8 DUF26 domains may be central to their ability to interact with other proteins but also with potential ligands (Figure S13c). In case of PDLP5, the higher ω values on the surface could indicate fast evolution events leading to sub- or neofunctionalization, as the PDLP5 orthologs all originate from the more recent duplication in the lineage leading to Brassicaceae species. The drastically different surface charge properties of related PDLPs from the same species (Figure 5b) suggest that different PDLPs and other DUF26 domain-containing proteins sense a rather diverse set of ligands. While the nature of these molecules is currently unknown, cell-wall derived carbohydrates or small extracellular molecules represent candidate ligands, but we were not able to identify any in our experiments (Figure S13a-b). Notably, we observed typical lectin-dimers in our PDLP5 and PDLP8 crystals, in which two lectin domains dimerized along an extended anti-parallel β-sheet (Figure 5c)^47^. In principle, this mode of dimerization could give rise to an extended binding cleft for a carbohydrate polymer, and presents an attractive receptor activation mechanism for PDLPs and CRKs, in which a monomeric ground state forms ligand-induced oligomers, as previously seen with LysM-domain containing carbohydrate receptors in plants^48^.

**Figure 5:**
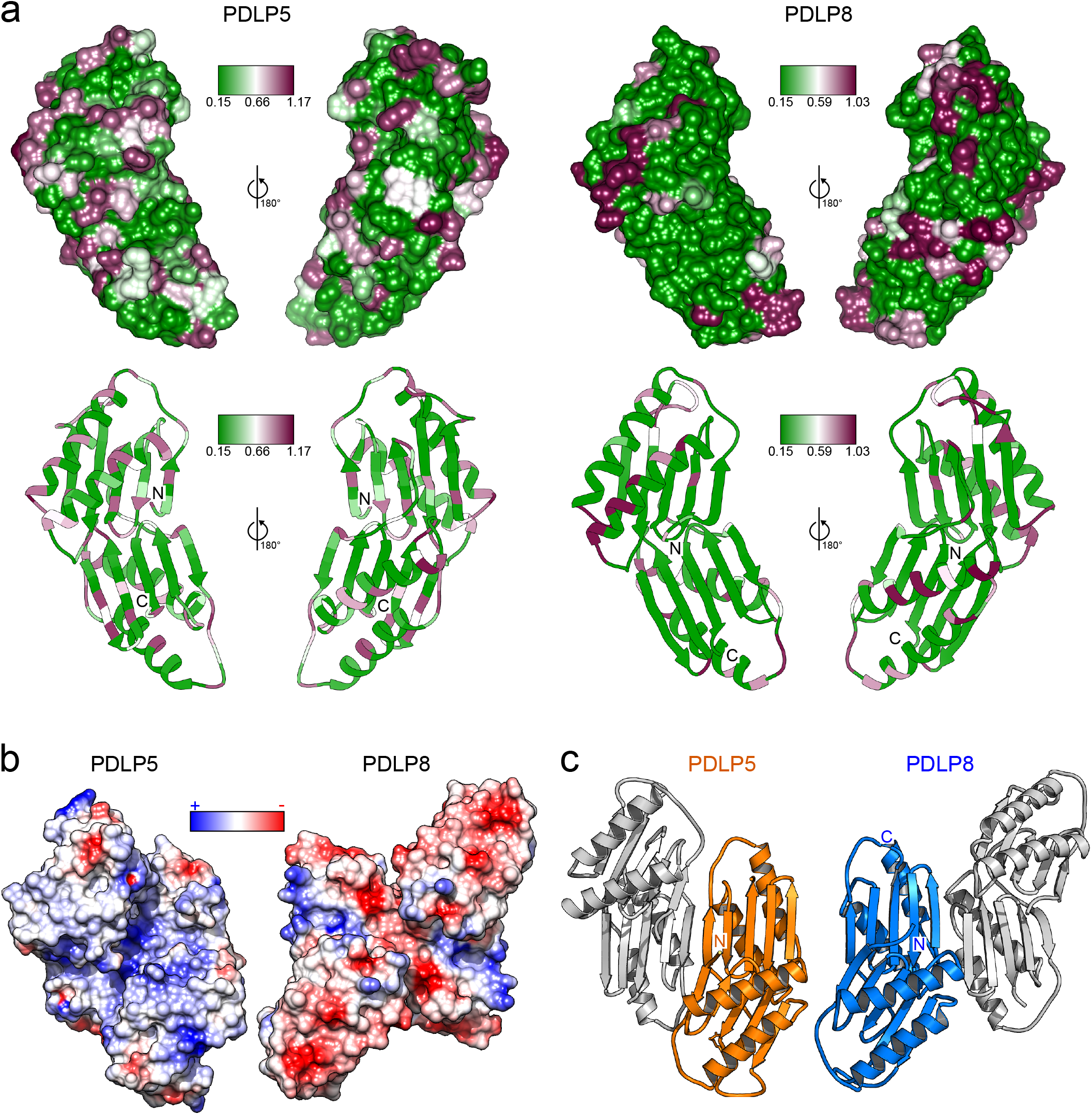
PDLP5 and PDLP8 may have drastically different oligomerisation modes, surface charge distributions and surface exposed residues are not widely conserved. **a)** The conservation of amino acid residues illustrated on the molecular surface of the PDLP5 or PDPL8 crystallization dimers, respectively. Site-wise ω (dN/dS) values, indicating the intensity and direction of selection on amino acid changing mutations, illustrated on the molecular surfaces (upper) and in ribbon diagrams (lower) of PDLP5 or PDPL8. The ω values range from 0.15 (green) to slightly over 1.0 (magenta), reflecting conserved sites under purifying selection and sites evolving close a neutral process, respectively. **b)** Electrostatic potential mapped onto molecular surfaces of the putative PDLP5 and PDLP8, orientation as in c) dimer, respectively. **c)** Ribbon diagrams of PDLP5 (orange) and PDLP8 (blue) crystallographic dimers. In both dimers large, antiparallel β-sheets are formed, using different protein-protein interaction surfaces.

### The CRK kinase domain is related to LRR and S-locus RLKs

Kinase domains transduce signals by phosphorylating substrate proteins and thereby are determining factors for signal specificity. Typically, the intracellular kinase domain has been used to investigate phylogenetic relationships between RLKs^9,15,16^. The typical CRK kinase domain is similar to the kinase domain of S-locus lectin and LRR RLKs from LRR_clade_3^15^ (Table S3). Based on the sequence of catalytic motifs in kinase domains^49^ most CRKs seem to be active protein kinases and the *in vitro* activity of several CRKs has been experimentally confirmed^25,28,30^. Most CRKs belong to the RD type^50,51^ which is considered to be capable of auto-activation but a few non-RD CRKs are present in plant genomes^49^.

Analyzing ectodomains and kinase domains of CRKs separately suggests that *Selaginella* ddCRKs share an ancestor with bCRKs, while *Selaginella* sdCRKs share an ancestor with vCRKs (Figure 6a). The clear separation of DUF26-A and DUF26-B (Figure 2d) and the timing of those events does not reveal whether the duplication of the DUF26 domain in the extracellular region of CRKs has happened more than once or whether functional constraints in the kinase domain led to the conserved similarity of *Selaginella* sdCRKs and vCRKs. Juxtaposition of phylogenetic trees based on ectodomains and kinase domain suggests several exchanges of kinase or extracellular regions among CRKs during evolution (Figure 6a). Most strikingly, a group of monocot-specific CRKs separates from other CRKs in a phylogenetic tree based on the kinase domain (Figure 6a). Those CRKs have an atypical gene model comprising a kinase domain with high similarity to concanavalin A-like lectin protein kinase domains (Table S3), and a different exon-intron structure (Figure 6b, S1b), altogether suggestive of chimeric gene formation following a tandem duplication^52^. The switch of the kinase domain and the associated changes in exon-intron structure is specific to grasses (*Poaceae*) and has likely resulted in a different set of target substrates. Exchange of kinase domains is not the only alteration of domain composition within DUF26-containing proteins. Loss of ectodomains and TMRs has established CRCKs at least three times; one group of CRCKs is specific to angiosperms (CRCK-I clade), one is specific to *Brassicaceae* and one only to *Arabidopsis thaliana*.

**Figure 6.**
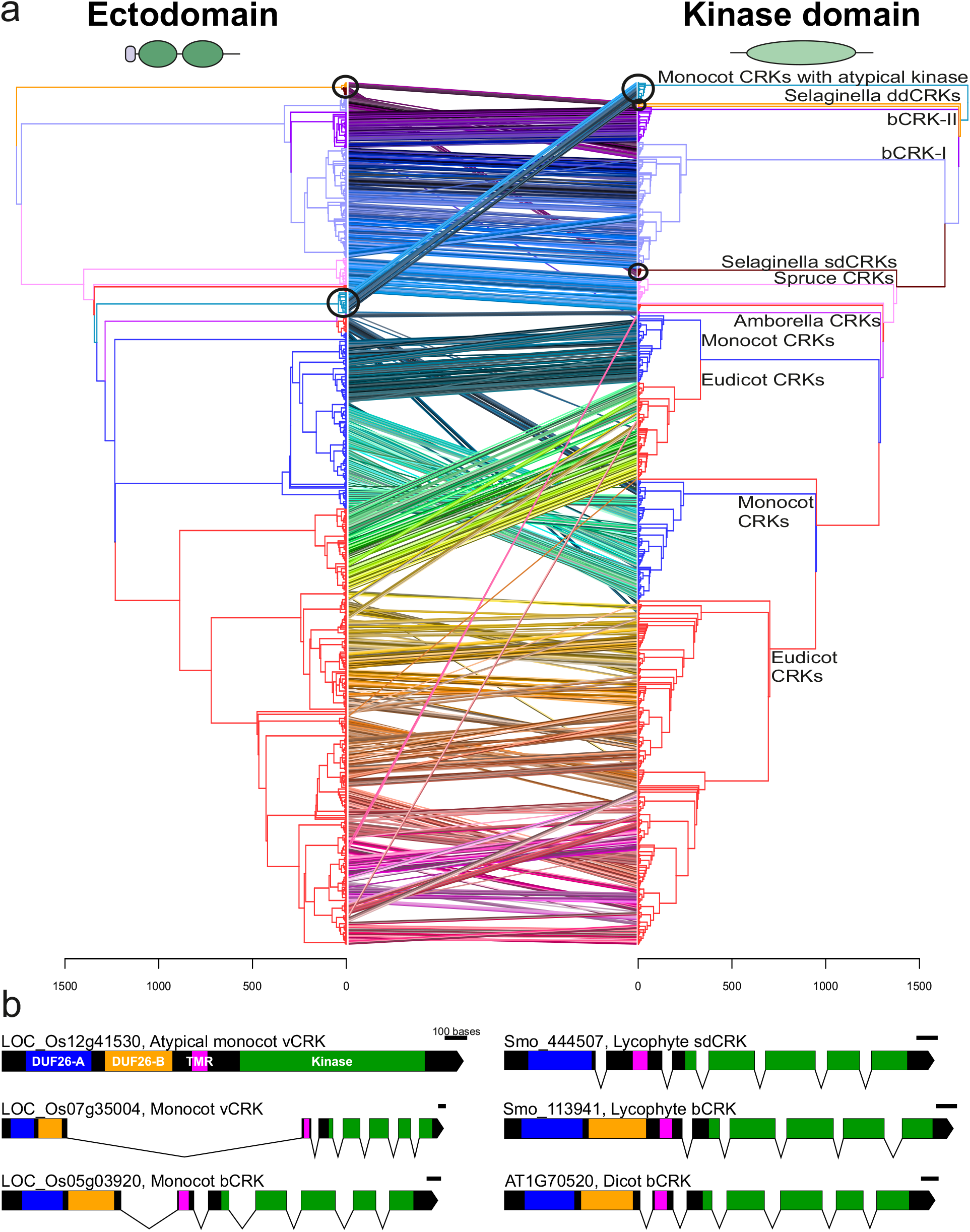
CRKs experienced domain rearrangements. **a)** Comparison of phylogenetic trees based on ectodomain region and kinase domain of 880 CRKs. Phylogenetic maximum-likelihood trees are presented as tanglegram where the tree of the CRK ectodomain region is plotted against the tree of the kinase domain. The kinase tree is rooted to atypical monocot CRKs with a Concanavalin-A type kinase domain and the ectodomain tree is rooted to CRKs from *Selaginella moellendorffii*. The ectodomain tree was detangled based on the kinase domain tree. Lines connect the ectodomain and kinase domain belonging to same gene, and connection are drawn in different colors for better visibility. Juxtaposition of the trees shows rearrangements and domain swaps of ecto- and kinase domains. Black circles highlight the difference between the ectodomains and kinase domains of the *Selaginella* sdCRKs and ddCRKs and also the group of the atypical monocot CRKs which have exchanged the kinase domain. **b)** The exon-intron structure of the CRKs. Usually CRKs contain seven exons: one encoding DUF26 domains, one encoding transmembrane region (TMR) and five exons encoding the kinase domain. In atypical monocot CRKs with exchanged kinase domain, whole gene is encoded by one or two exons. The scale bar for each gene represents 100 bases. Regions encoding the DUF26-A are colored with blue, the DUF26-B with orange, the transmembrane region (TMR) with pink and the kinase domain with green.

### Mixed-mode evolution of large gene families

In order to carry out more detailed analyses of gene family dynamics we analyzed the synteny, conservation of the gene order between species, as well as tandem duplications in highly contiguous chromosome-level assemblies of *Amborella trichopoda*, tomato (*Solanum lycopersicum*), Arabidopsis, rice and maize (*Zea mays;* Figures 7a and S7), and estimated the timing of the duplication events by reconciliation of gene trees with species trees (Figure S14a).

**Figure 7.**
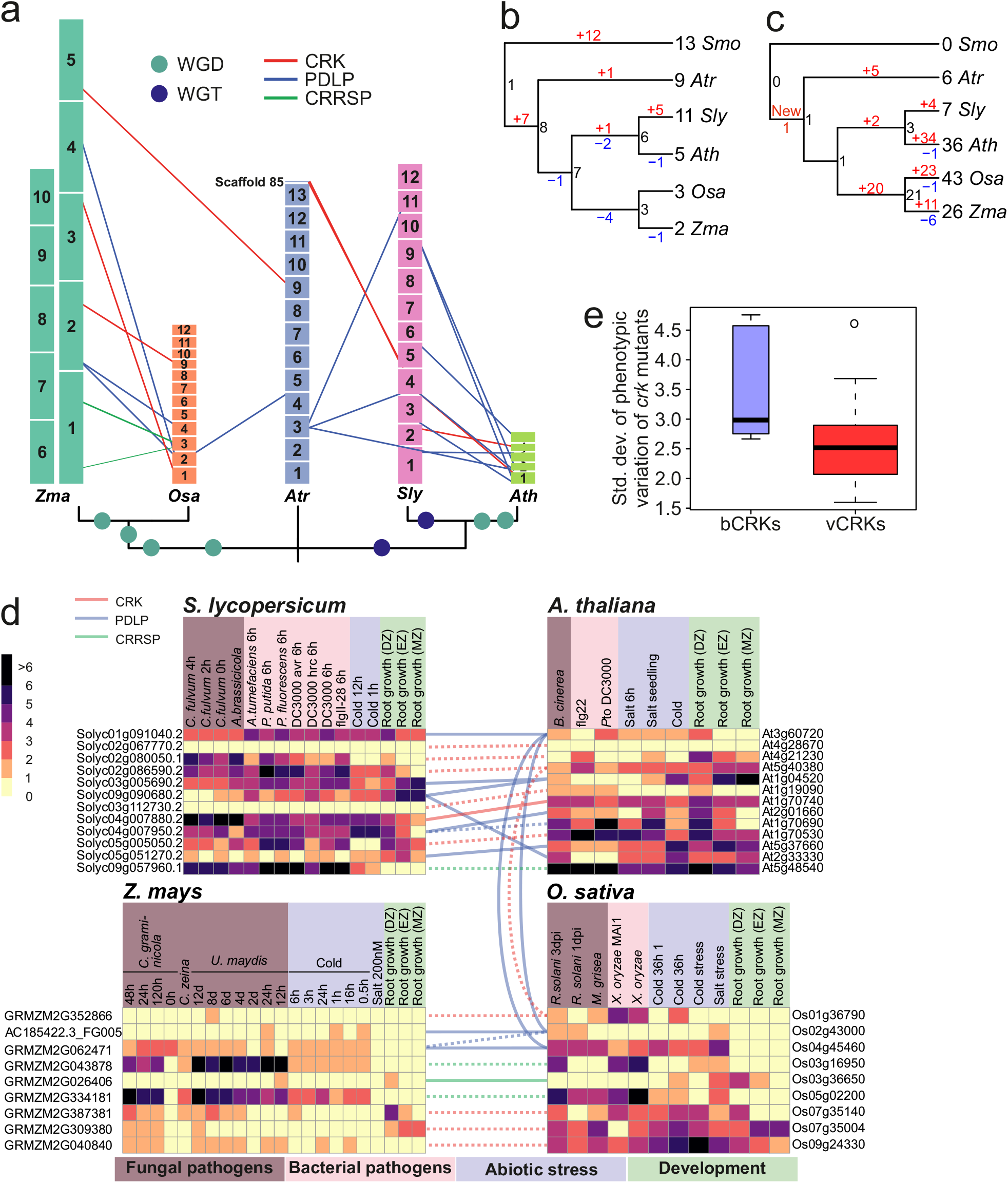
Identification of the modes of gene family evolution in DUF26-containing genes in *Arabidopsis thaliana*, tomato, rice, maize and *Amborella trichopoda*. **a)** Gene families that are preferentially retained after whole genome multiplications (WGMs) are typically identified by synteny analysis. The figure illustrates syntenic regions containing DUF26 genes from *Amborella trichopoda* to monocots *Oryza sativa* and *Zea mays* (to left from middle) and to eudicots *Solanum lycopersicum* and *Arabidopsis thaliana* (right from the middle). In the synteny analysis within monocots and dicots, segments with at least 5 syntenic genes were included, whereas in comparisons to *Amborella* the minimum threshold was 3 syntenic genes. Analyses were carried out with Synmap software within CoGe. For *Amborella trichopoda* genomic locations of DUF26-containing genes are only known on chromosome/scaffold level based on physical mapping. **b and c)** Gene families with a preferential retention pattern after WGMs show conserved gene counts over species. Phylogenetic tree of the five species shown in the panel was used to reconcile the gene trees and estimate gene counts in ancestral nodes for **b)** bCRKs and **c)** vCRKs, using *Selaginella moellendorffii* as outgroup. The gains are highlighted with red and losses with blue. **d)** Gene families with preferential retention pattern should have many orthologs. Heatmaps of the normalized transcriptional expression counts (Transcript per million [TPM]) of candidate DUF26 orthologs from four of the species: *Solanum lycopersicum*, *Arabidopsis thaliana*, *Zea mays*, and *Oryza sativa*. Coloring in heatmaps is proportional to log_2_ (TPM) value that represents the gene expression level. The corresponding log_2_ (TPM) value is displayed next to the color key. The rows represent gene models and the columns show the experiments, collected from publicly available Sequence Read Archive (SRA) database. SRA accessions are annotated to relevant stress conditions (descriptions are presented in Table S4). Solid lines connect putative orthologs based on evidence from phylogenetic and synteny analyses; dashed lines connect putative orthologs based on evidence from either phylogenetic or synteny analyses. **e)** Final prediction of gene families evolving under dosage balance is that their knockouts demonstrate a high phenotypic effect. This can be seen by reanalysis of phenotype data from (Bourdais *et al*.^24^); the bCRK T-DNA insertion mutants display a significantly larger standard deviation (Y-axis) over different phenotyping experiments than vCRK mutants. Pathogens: *Agrobacterium tumefaciens*, *Alternaria brassicicola*, *Botrytis cinerea*, *Cercospora zeina*, *Cladiosporum fulvum*, *Colleotrichum graminicola*, *Magnaporthe grisei*, *Pseudomonas putida*, *Pseudomonas fluorescens*, *Pseudomonas syringae pv. tomato DC3000*, *Rizoctonia solani*, *Ustilago maydis*, *Xanthomonas oryzae*.

Within the young, rapidly diverging β-group the vCRKs show large lineage-specific expansions. The ancestral origins for monocot and eudicot vCRKs differ, and neither synteny nor orthology can be identified (Figure 7c and S14c). Altogether this suggests that this younger subfamily has a high birthrate and that it expands rapidly by tandem duplications in all species. Additionally, many of the tandems are lost or fractionated after WGMs. Similarly the CRRSPs demonstrate little synteny between different species (Figure 7a and S14d), and CRRSPs in rice and Arabidopsis experienced lineage-specific tandem duplications (Figure S14d). In *Brassicaceae* this expansion can be traced to *Amborella* CRRSP (*AtrCRRSP2*), altogether suggesting a tandem mode of expansion.

Tandem duplications evolve through unequal crossover or homologous recombination events^53^. Unequal crossover produces copy number variation, whereas homologous recombination such as gene conversion plays a role in concerted evolution, which can maintain the similarity between gene copies over long periods^54^. Gene conversion is known to depend on the genomic distance as well as sequence homology. Accordingly, we observed several events among the lineage-specific tandem vCRK expansions (Table S4), whereas in case of bCRKs, events were observed only in the tandem expansion in *Amborella*. This suggests that gene conversion is an important process maintaining the similarity between recent tandem duplicates but as the sequences diverge over time the conversion events become increasingly rare.

The CRCK-I genes are present in most genomes as single copy genes within conserved syntenic genome segments, suggesting that duplicates from WGD events have been lost during genome fractionation (Figure 7a). The evolution follows a specific dosage balance model where the maintenance of a single copy is critical to the organism.

A hallmark for gene families evolving under dosage balance is that their overall numbers should be conserved among species with similar WGM history. In the species tree (Figure 7a), most of the branches contained one or two WGMs. Despite these events, the overall number of bCRKs is well conserved in angiosperms (Figure 3b, 7b S3b and S7b). However, in *Amborella trichopoda* five bCRK genes appear in tandem and these genes are at the roots of the respective orthologs (Figure S6b), indicating an ancestral SSD origin still present in *Amborella*. The duplicate region experienced considerable fractionation during evolution leading to *Brassicaceae* and *Solanaceae* lineages, resulting in scattered bCRK-I orthologs with little conserved synteny, whereas in the two grasses the tandem duplicate was lost altogether. This indicates rapid pseudogenization of the duplicated tandem blocks after WGMs, with, except for *Solanaceae*, no recent tandem expansions. Altogether this suggests that a gene family that initially existed as a tandem duplicate may have shifted towards a dosage balance mode of evolution. Dosage balance is observed in the second subfamily of ancient origin, PDLPs, since they appear in genomic regions where synteny is conserved within eudicots and monocots (Figure 7a), and no recent or ancient SSD events can be detected.

One of the predictions for the gene families evolving under dosage balance is that retained duplicates should exhibit less functional divergence than other duplicates^3^. We explored functional conservation by analyzing publicly available gene expression data on stress treatments (Table S5; Figure 7d, S15). In agreement with studies in Arabidopsis and rice^24,26,31,55^, pathogen treatments have the biggest impact on transcript abundance of DUF26-containing genes, in particular CRKs and CRRSPs (Figure S15). Our analysis of gene expression data suggests extensive lineage-specific functional diversification. This is visible in the correlation rank between putative orthologs; in many cases higher correlation can be found with DUF26-containing genes that have less similarity in sequence, indicating that the closely related genes have undergone sub- or neo-functionalization following duplications^56,57^.

Overall bCRKs show elevated transcript levels in response to stress treatments, while many vCRKs have elevated transcript levels in pathogen treated samples. Rice PDLPs display altered transcript levels in some specific stress treatments. Despite re-arrangements and lineage-specific expansions the data provide support for seven putative orthologs, including three PDLP and three CRRSP relationships (Figures 7d; Table S5). Even though the synteny of bCRKs (and PDLPs) is more conserved compared to CRRSPs, bCRKs demonstrate varying responses to stimuli, whereas in CRRSPs synteny is associated with similar functions.

The second prediction from the dosage balance model is that since the protein products of the genes are highly connected and thus interact with many other proteins, disturbances in the dosage balance should have large effects on an organism’s phenotype^58^. Reanalysis of phenotyping data of T-DNA mutant insertion lines^24^ confirms that bCRKs indeed demonstrate a larger variance in phenotypes than vCRKs (p=0.03; Wilcox test; Figure 7e). Altogether the analysis suggests that PDLPs and bCRKs are evolving according to the dosage balance model, whereas the vCRKs and CRRSPs evolve by SSD mechanisms.

## Discussion

Compared to animal genomes, plant genomes encode a large number of large gene families^59^. In particular signal transduction components including transcription factors, protein kinases and phosphatases have experienced drastic expansions in plants^59^. This might reflect the adaptation to a sessile lifestyle but also could indicate a different strategy for signal transduction and integration at the cellular level. The large, in part lineage-specific expansions and conversions between different domain arrangements seriously hamper the identification of orthologous proteins in different plant species. Here we studied the evolution of a large plant protein family which is hallmarked by heterogenous domain architecture and drastic lineage-specific expansions of subgroups, the DUF26-containing proteins. We identified 1409 high-quality gene models representing CRRSPs, CRKs and PDLPs from major plant lineages. Our analyses suggest that sdCRRSPs are the ancestral type of DUF26-containing proteins. CRKs originated from a fusion of CRRSPs with TMR and kinase domain of LRR_clade_3 RLKs^15^ in the lineage leading to lycophytes. PDLPs and ddCRRSPs emerged subsequently through the loss of the kinase domain or the TMR and kinase domain. Our results reveal an ancient split into two distinct groups. The α-group is strongly conserved in size and sequence throughout embryophytes. This facilitates identification of functional orthologs and extrapolation of functional information from model plant species to crops. The β-group evolved before the split of monocots and eudicots and contains CRKs and CRRSPs that expanded through WGDs followed by lineage-specific tandem duplications. Domain re-arrangements in the β-clade led to secondary groups of ddCRRSPs and sdCRRSPs while the recruitment of a different kinase domain in grasses suggests the re-routing of signaling pathways towards novel phosphorylation substrates. Thus, it is likely that members of the β-group have been subject to sub- and neo-functionalization, which is a challenge for functional analyses. The domain exchanges in DUF26-containing proteins highlight the importance of comparative analysis of phylogenies inferred from full-length protein sequences with those inferred from individual domains. WGDs have been associated with periods of environmental upheaval and an increase in biological complexity^2,60^. Accordingly, the appearance and radiation of DUF26-containing proteins with different domain structures as well as CRK and CRRSP expansions co-occur with the evolution of novel physiological characteristics, such as vasculature, and with the adaptation to new habitats and lifestyles (Figure 1b).

Sequence analysis suggested that DUF26 proteins could be specific to embryophytes. Crystallographic analysis of two PDLP ectodomains reveals that the structure of the DUF26 domains closely matches the fold of the sdCRRSP Gnk-2, which is evolutionarily distant from the PDLPs. PDLPs contain two DUF26 domain and the structure of Gnk-2 is more similar to the DUF26-A. However, despite the high structural similarity the mannose-binding function of Gnk-2 is not conserved in the PDLP DUF26-A domain. Intriguingly, plant DUF26 domains share significant structural similarity to fungal carbohydrate-binding modules. Notably, the tandem arrangement of two lectin-like DUF26 domains appears to be plant-specific. Rapid sequence divergence^61^ is a limiting factor in detection of homology at the amino acid sequence level, seen e.g. in the marked differences between DUF26 from *Marchantia polymorpha* and *Physcomitrella patens* and those from other plants. This may obscure identification of DUF26 domains in charophytes and other algal species. The physiological ligands of ddCRRSP, CRKs and PDLPs remain to be discovered and our work suggests that different tandem DUF26 domains likely recognize diverse sets of ligands. Similar to plant malectin receptors^62^, DUF26 domains may have evolved novel or additional functions which might include mediation of protein-protein interactions at the cell surface^20,35^. The strong structural similarity between DUF26 domains and fungal lectins suggests a common origin, and DUF26 proteins represent novel carbohydrate-binding domains in plants. Identification of ligands for different DUF26-domains will provide novel insights in to perception of cell wall status or environmental signals. However, this may be challenging since plant cells and their cell walls contain a large number of carbohydrates and related compounds.

From the evolutionary analysis, an overall model emerges (Figure 8). The young gene families initially expand through tandem duplications and therefore experience more relaxed selection^63^. This is supported by the fact that the tandem genes function in processes that require fast responses such as adaptation to environment, pathogen responses and secondary metabolism^2,64^, and that these gene families show high variation across species and have high kn/ks rates^5^. In tandems, main evolutionary forces are unequal crossover and concerted evolution through gene conversion, but over time the genes evolve into their specific functions. This process may be interrupted by WGM events. Since the tandem genes are not evolving under dosage balance, there is no compensatory drift^65^, and thus drift and selection by dosage eventually drives one of the duplicates into fixation while others turn into pseudogenes. Assuming that the elements driving tandem duplications are still present after fractionation, the remaining duplicates may in turn expand. In case of a tandem where all genes have established a unique functional role in the system, drift may drive the duplicated tandem into scattered orthologs. These orthologs may eventually assume a fixed syntenic position in the genome and switch to a dosage balance mode of evolution. The evolutionary mode of the gene family would depend on the balance between the death rate after WGMs and the birth rate of the tandem duplications.

**Figure 8.**
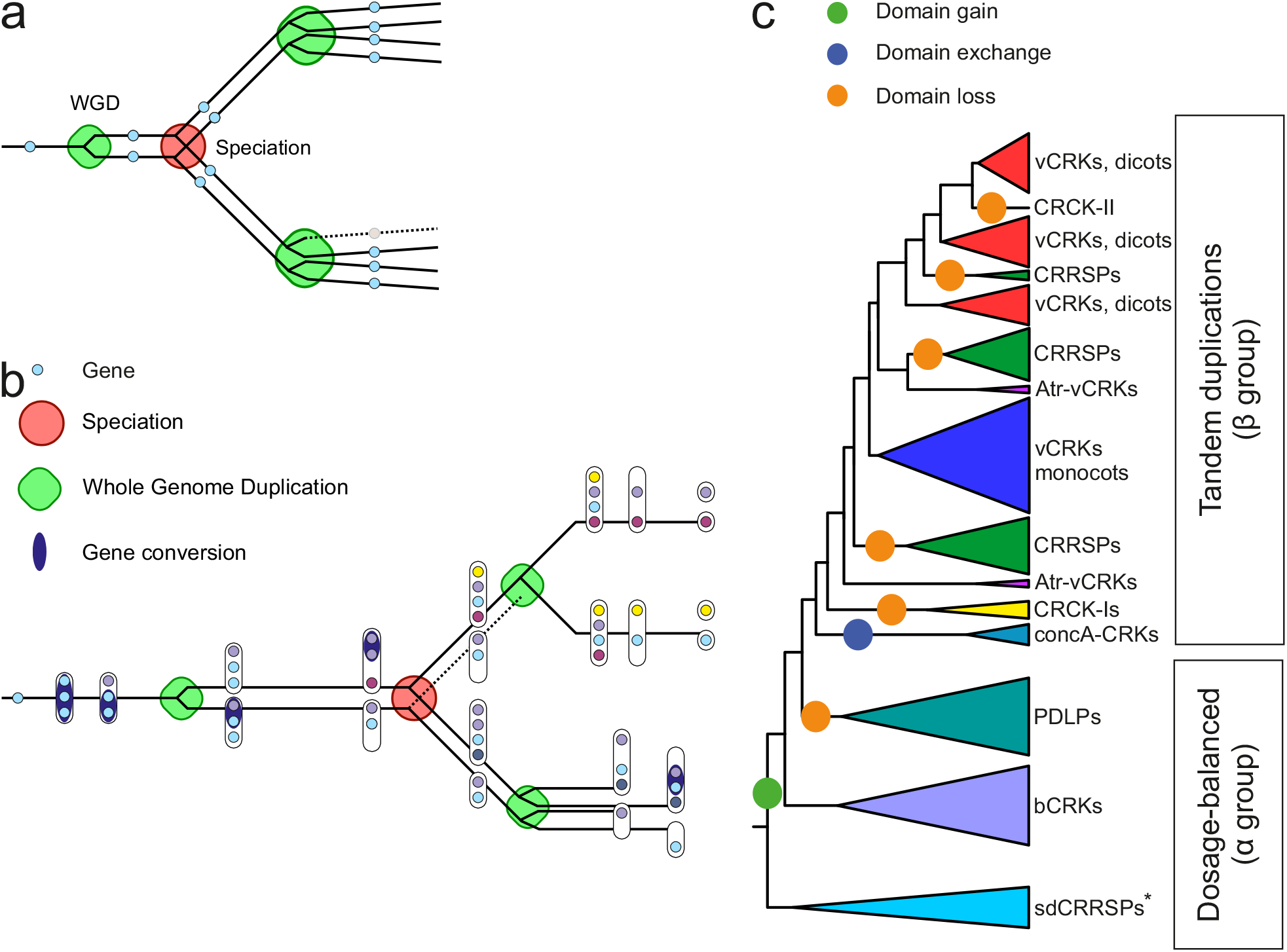
Model of mixed-type gene family evolution. Gene families evolve through two major events, whole genome multiplications (WGM) and small-scale duplications (SSD). Genes related to environmental responses and secondary metabolism experience SSDs in the form of tandems, whereas highly connected genes associated with transcriptional and developmental regulation or signal transduction functions are preferentially retained after WGMs. **a)** Prevailing hypothesis for the retention pattern is dosage-balance; in case of highly connected genes the stoichiometric balance needs to be maintained, and therefore selection acts against gene losses after WGMs and against duplications by SSDs. **b)** On the other hand, gene family evolving through tandem duplications **(b**; evolution before the speciation node) has a high birth rate and therefore the number of duplicates between species can vary. After duplications the homogeneity of the duplicates is maintained through gene conversion events, which has a high probability with near-by homologous sequences. This can be maintained for long periods, but eventually over time the sequences diverge by drift and selection based on dosage. Our data suggests that a tandemly expanding gene family may evolve into a dosage balance mode as a result of WGMs **(b**; evolution after speciation node). Following WGMs, the duplicated tandems may experience extensive fractionation due to drift and selection by dosage which fragments the tandem stucture. At the same time, the connectivity of the gene family has been accumulating through sub- and neofunctionalization, increasing pressure for retention of the gene models. These phenomena together may result into a dosage balance model of evolution (top branch after speciation node). This does not necessarily occur across all WGM events and depends on the tandem duplication rate, as was observed for bCRKs in Solanaceae (bottom branch), where there exist both single copies and a later tandem expansion in the genome. Different subfamilies can be in different states of this process. **c)** CRRSPs and PDLPs follow dosage balance mode after the paleohexaploid event, whereas bCRKs have assumed the mode in later WGM events. The overall numbers of the bCRKs are preserved but identification of orthologs between species that have experienced independent WGMs is difficult, suggesting that convergent functionality of the members is recent. Gene families expanding through tandem duplications such as vCRKs and CRRSPs have high birthrate and demonstrate several lineage-specific expansions. * Lost in Brassicaceae species and rice

Our study of DUF26-containing proteins demonstrates both the challenges of analyses of large protein families and the power of combining advanced evolutionary and structural methods. Our analysis will provide a model for future studies of similarly large protein families and will facilitate the forthcoming detailed biochemical and physiological investigation of the mechanistic functions of CRKs, PDLPs and CRRSPs in different plant species.

## Materials and methods

### Gene identification and annotation

Altogether 32 plant and algae genomes (Table S1) covering the major plant lineages were selected for analyses. For 27 species protein annotations (primary transcripts) and genome sequence data was retrieved from Phytozome^66^, and Barley (*Hordeum vulgare*) from Gramene (http://www.gramene.org) with the latest names for gene models from IPK server (http://webblast.ipk-gatersleben.de/barley_ibsc/)^67^. Silver birch (*Betula pendula*) was sequenced at the University of Helsinki^56^. Eggplant (*Solanum melongena*) data was retrieved from Eggplant Genome DataBase (http://eggplant.kazusa.or.jp/). *Klebsormidium flaccium* and Sacred lotus (*Nelumbo nucifera*) genome data were from NCBI (https://www.ncbi.nlm.nih.gov). Additionally the FungiDB^68^ (www.fungidb.org), InsectBase^69^ (http://www.insect-genome.com) and human (*Homo sapiens*), chicken (*Gallus gallus*) and zebrafish (*Danio rerio*) genomes were screened for DUF26. Detailed information of the genome versions and references are given in the Table S6.

HMMER (version 3.1b2) search^70^ for PFAM domain with ID PF01657 (stress-antifungal domain) was carried out among AA sequences representing gene models from different species^71^. Genome sequences were checked with Wise2 (version 2.4.1) software^72,73^. All gene models found with HMMER were manually curated, and new genes found with Wise2 were manually annotated using Fgenesh+^74^. Birch (*Betula pendula*)^56^ and Sacred lotus (*Nelumbo nucifera*) were fully manually annotated as they did not have gene models *a priori*. High rates of manual annotation and curation were needed for *Selaginella moellendorffii*, grapevine (*Vitis vinifera;* version Genoscope.12X^75^) and potato (*Solanum tuberosum*). Sequences from each species were further checked by carrying out a multiple sequence alignment and phylogenetic tree estimation with PASTA^76^. Partial gene models were identified by checking sequences individually. Genes were defined as pseudogenes if the genomic sequence was available but no full domain structure could be predicted. In cases where the prediction problem was caused by the length of the contig or a gap in the genome sequence the gene model was marked as partial. Pseudogenes and partial gene models were not included in the subsequent analyses.

For domain analyses and phylogenetic trees containing only domain sequences, the domain borders were defined with HMMER using the PFAM domain PF01657 for DUF26, and PF07714 for the kinase domain from curated dataset. The ectodomain region was defined to end at the border of the transmembrane region in the PDLPs and CRKs. The partial PDLP from *Marsilea quandrifolia* was identified by using pBLAST search against sequences in the NCBI database.

### Phylogenetic trees

Only full gene models were used to infer phylogenetic trees. Sequence quality in alignments was checked using Guidance (version 2.01) and alignments were built using the MAFFT option^77^. Sequences with low quality score were removed from datasets and alignments were built again with PASTA. For phylogenetic trees, alignments were filtered in in Wasabi^78^ to remove residues with less than 10 percent coverage. Filtering was required due to the high sequence diversity (on less conserved regions) resulting in a high number of gaps in multiple sequence alignments. Maximum likelihood (ML) phylogenetic trees were inferred for filtered and also unfiltered data using RAxML (version 8.1.3)^79^.

ML phylogenetic trees were bootstrapped using RAxML (version 8.1.3) for 1000 bootstrap replicates. For phylogenetic trees containing full length sequences with all domain structures bootstrapping was also carried out with partitioning (both DUF26 and kinase domains defined separately). The PROTGAMMAJTT model was used in phylogenetic analyses using RAxML. Model selection was based on a Perl script for identifying the optimal protein substitution model (available in RAxML webpage, provided by Alexandros Stamatakis). Bootstrapped trees are available on Wasabi^78^ (see figure legends). Comparison on phylogenetic trees based on CRK ectodomain and kinase domain regions was visualized in R using the “dendextend” package.

### Exon intron structure

The number of exons for all genes was estimated using Scipio (version 1.4.1)^80^ using default parameters (minimum identity of 90% and coverage of 60%). It internally uses BLAT to perform the initial alignment of the protein sequences against the genome followed by refinement of hits to determine the exact splicing borders and to obtain the final gene structure. The number of exons per gene was extracted from the final result.

### Orthogroup generation

11 representative species from different clades (*Arabidopsis thaliana*, *Amborella trichopoda*, *Oryza sativa*, *Zea Mays*, *Vitis vinifera*, *Populus trichocarpa*, *Aquilegia coerulea*, *Brachypodium distachyon*, *Physcomitrella patens*, *Selaginella moellendorffii* and *Spirodela polyrhiza*) were chosen to study the evolution of the DUF26-containing proteins. Primary protein sequences of these 11 species were downloaded from Phytozome (version 11.0). An all-against-all BLAST was run for all the protein sequences followed by generation of orthogroups using the software OrthoMCL (version 2.0.9)^81^ with an inflation parameter of 1.5 for the clustering phase. Clustering yielded 34,535 orthogroups.

### Species tree generation

Orthogroups containing one representative protein for each of the 11 species were chosen to generate the species tree. Multiple sequence alignment was carried out on the single copy orthogroups using PRANK^82^ and the output was used to infer a species tree using RAxML^79^.

### Evolutionary rate and ancestral size estimation

The evolutionary rate and ancestral size of the orthogroups were modelled using Badirate software (version 1.35)^41^. The species tree and orthogroups generated from the previous steps were used as input for Badirate. The BDI (Birth, Death, Innovation) rate model was used. The Free Rates (FR) branch model was chosen which would assume every branch of the species tree to have its own turnover rates. Turnover rates of orthogroups were estimated using the maximum likelihood fitting. Orthogroups were defined as protein kinases if they included sequences with PFAM domain PF00069. Orthogroups containing RLKs were defined based on known *Arabidopsis* RLKs^15^. Plasmodesmata-related orthogroups were defined based on *Arabidopsis thaliana* genes related to plasmodesmata^40^.

### Nucleotide CDS sequence generation from protein sequence for PAML

The GFF file output from Scipio^80^ was pre-processed by an in-house script and processed with the gff3 module of the GenomeTools (version 1.5.4)^83^ software. The final GFF file along with the corresponding species genome in fasta formatted file was passed as an input to the extractfeat module of the GenomeTools software to extract the final nucleotide CDS sequences.

### PAML analyses

We estimated *d*_*N*_/*d*_*S*_ ratios (ratio of non-synonymous and synonymous sites, ω) for conserved clades (bCRK-I, bCRK-II, CRCKs (orthologs of AtCRK43), PDLPs and sdCRRSPs) from eleven species (*Arabidopsis thaliana*, *Amborella trichopoda*, *Oryza sativa*, *Zea Mays*, *Vitis vinifera*, *Populus trichocarpa*, *Aquilegia coerulea*, *Brachypodium distachyon*, *Physcomitrella patens*, *Selaginella moellendorffii* and *Spirodela polyrhiza*) by using the codeml program from PAML (version 4.9)^84^. We applied the one-ratio model (M0) to estimate overall *d*_*N*_/*d*_*S*_ ratios for each conserved group separately and free ratios neutral model (M1) to estimate *d_N_*/*d_S_* ratios for each branch within conserved clades^85^. To study the evolution of PDLP5 and PDLP8, sitewise-analyses of their homologs was carried out. As PDLP5 is specific to *Brassicaceae*, we additional nucleotide sequences for orthologs of AtPDLP5 from NCBI, Phytozome and CoGe databases. Furthermore, additional sequences for orthologs of AtPDLP8 were included in the alignment to improve depth and reliability of the analysis. Multiple sequence alignments of coding nucleotide sequences were constructed with PRANK^82^ and phylogenetic trees were estimated using RAxML^79^ for codeml.

### Syntenic vs tandem duplications

Syntenic and tandem duplications were analysed using Synmap application in CoGe^86^, using default settings. Tandem duplications were defined as genome regions with at least three to five duplicate genes (Table S4). Synteny comparisons were done between *Arabidopsis thaliana* and *Solanum lycopersicum*, *S. lycopersicum* and *Amborella trichopoda*, *A. trichopoda* and *Oryza sativa* and *Zea mays* and *Oryza sativa*. Tandem duplication results from DAGchainer were collected for each species. The results were filtered based on annotated gene models from selected species. The currently available *Amborella trichopoda* genome is presented only as scaffolds, and the genes were placed to chromosomes based on physical mapping^87^. Scaffolds not assigned to any chromosome were added separately. Thus the location of the *Amborella trichopoda* genes in the genome is only a rough estimate (Figure 7a).

### Gene conversion analyses

Gene conversion events were estimated from nucleotide sequences for the same eleven species that were analyzed for *d*_*N*_/*d*_*S*_ ratios with GENECONV (version 1.81a)^88^. Analyses were carried out for the main clades of the eleven species. Tor bCRKs and vCRKs separate analyses were carried out using sequences from the five species used in synteny analyses (*Arabidopsis thaliana*, *Amborella trichopoda*, *Oryza sativa*, *Solanum lycopersicum* and *Zea mays*). The largest tandem region of vCRKs in *A. thaliana* chromosome 4 was analyzed separately to validate the results from the analysis with all vCRKs from *A. thaliana*.

### Gene tree reconciliation

Gene tree reconciliation was carried out using DLCpar (version 1.0)^89^ downloaded from https://www.cs.hmc.edu/~yjw/software/dlcpar/. NCBI taxonomy was used as the species tree, downloaded in newick format from PhyloT website, http://phylot.biobyte.de/. Reconciliation was carried out using DLCpar search with 20 prescreening iterations, followed by 1000 search iterations. The solution was visualized in R, using custom scripts and ‘ape’ package.

### Phenomics data analysis

Phenotyping data of T-DNA mutant insertion lines was normalized against the Col-0 data by calculating Z-scores, see Bourdais *et al*.^24^ The standard deviation (SD) over all experiments was calculated for each allele, and in case of several insertion alleles the one with maximum SD was selected. The residuals of the bCRK vs vCRK split in the data were tested for normality using Shapiro’s test. Since the null hypothesis (normality) was rejected with p<0.05 the difference between groups was tested with Wilcox test.

### Transcriptomic analyses

Paired end RNAseq data was collected from the publicly available sequence read archive (SRA) database by fastq-dump.2 (version 2.5.7) for *Arabidopsis thaliana*, *Oryza sativa*, *Solanum lycopersicum* and *Zea mays*. FastQC (version 0.11.4) (https://www.bioinformatics.babraham.ac.uk/projects/fastqc/) was used to check the quality of the samples. Low quality reads and bases were removed by Trimmomatic (version 0.36)^90^ with the following options: phred33, TRAILING: 20, and MINLEN: 30. Filtered reads were mapped to gene models from Phytozome version 12, by Kallisto, run in paired end mode (version 0.43.1, --bias and --bootstrap: 200)^91^. Bootstrap samples were averaged (custom R code) and gene expression abundance (transcript per million [TPM]) was estimated by tximport (version 1.2.0)^92^ followed by averaging over biological replicates. Ortholog comparison between species was carried out by grouping the experiments into seven categories, with maximum TPM among experiments representing gene response. Pearson correlation was calculated among orthologs and all other possible pairs.

### Protein expression and purification

An expression construct coding for the *PDLP5* ectodomain (amino acids 1-241) was codon optimized for *Spodoptera frugiperda* and synthesized by Geneart (Thermo Fisher). Using the PfuX7 polymerase^93^, the gene for the *PDLP8*-ECD (1-253) was amplified from *Arabidopsis thaliana* cDNA. The Gibson assembly method^94^ was employed to insert the *PDLP5* and *PDLP8* ectodomain coding sequences into an adapted pFAST-BAC1 vector (Geneva Biotech), providing a C-terminal 2x-STREP-9xHIS tag. *PDLP5* point mutations (C101A, C148A and C191A) were then introduced as described^95^. Bacmids were generated by transforming the plasmids (confirmed by sequencing) into *Escherichia coli* DH10MultiBac (Geneva Biotech). Virus particles were created by transfecting (Profectin, AB Vector) the bacmids into *Spodoptera frugiperda* SF9 cells. For secreted protein production, *Trichoplusia ni* Tnao38 cells were infected with a viral multiplicity of 1, incubated for 3 days at 22 °C. The protein-containing supernatant was separated from the intact cells by centrifugation and subjected to Ni^2+^-affinity chromatography (HisTrap Excel; GE Healthcare) in buffer A (10 mM Hepes 7.5, 500 mM NaCl). Bound proteins eluted in buffer A supplemented with 500 mM imidazole. The elution fractions were pooled and further purified by StrepII-affinity purification (Strep-Tactin XT Superflow high capacity, IBA) in buffer B (20 mM Tris pH 8.0, 250 mM NaCl, 1 mM EDTA). The column was washed with 5-10 column volumes of buffer B and eluted in buffer B supplemented with 50 mM biotin. The C-terminal 2x-STREP-9xHIS tag was subsequently removed by adding tobacco etch virus (TEV)-protease to the StrepII elution in a 1:100 ratio for 16h at 4 °C. The 2x-STREP-9xHIS-tag and the HIS-tagged TEV-protease were then separated from the respective ectodomain by an additional Ni^2+^-affinity chromatography step (HisTrap Excel; GE Healthcare). Cleaved PDLP5, PDLP5^C101A^, PDLP5^C148A^, PDLP5^C191A^ and PDLP8 ectodomains were next subjected to preparative size exclusion chromatography using either a HiLoad 26/600 Superdex 200 pg (PDLP5 and PDLP8) or HiLoad 16/600 Superdex 200 pg (PDLP5^C101A^, PDLP5^C148A^ and PDLP5^C191A^) column, equilibrated in 20 mM sodium citrate pH 5.0 and 150 mM NaCl. Monomeric peak fractions were collected and concentrated using an Amicon Ultra (Milipore) filter device. The concentrated monomeric peak fractions of PDLP5, PDLP5^C101A^, PDLP5^C148A^ and PDLP5^C191A^ were additionally subjected to analytical size exclusion chromatography on a Superdex 200 Increase 10/300 GL column (GE healthcare) equilibrated in 20 mM citrate pH 5.0 and 150 mM NaCl.

### Thermostability assay

20 μl reactions consisted of either PDLP5, PDLP5^C101A^, PDLP5^C148A^ and PDLP5^C191A^ ectodomains at a concentration of 1.5 mg/ml in 20 mM citrate pH 5.0, 150 mM NaCl, 10x SYPRO Orange dye (Thermo Fisher), and were mixed in a 384-well ABI PRISM plate (Applied Biosystems). Using a 7900HT Fast RealTime PCR system SYPRO Orange fluorescence was measured. The reactions were initially incubated for 2 min at 25 °C and then the temperature was increased to 95 °C at a heating rate of 0.5 °C/min. Resulting melting curves were fitted with a Boltzman function using GraphPad Prism and the melting temperatures, T_m_, correspond to the first inflection point of the Boltzman fit.

### Isothermal titration calorimetry

ITC experiments were performed at 25°C using a Nano ITC (TA Instruments, New Castle, USA) with a 1.0 mL standard cell and a 250 μl titration syringe. The PDLP5 ectodomain was gelfiltrated into ITC buffer (20 mM sodium citrate pH 5.0, 150 mM NaCl) and all carbohydrates were resuspended into ITC buffer. The experiments were carried out by injecting 24 times 10 μl of D-+-Mannose (1 mM; Sigma), Pectic Galactan (2mg/ml; Megazyme), Rhamnogalacturonan (2mg/ml; Megazyme), Polygalacturonic Acid (2mg/ml; Megazyme), Cellohexaose (1 mM; Megazyme) or Arabinohexaose (1 mM; Megazyme) aliqots into PDLP5 (~100 μM) in the cell at 150 s intervals. ITC data for the D-+-Mannose experiment were corrected for the heat of dilution by subtracting the mixing enthalpies for titrant solution injections into protein free ITC buffer. Data were analyzed using the NanoAnalyze program (version3.5) as provided by the manufacturer.

### Protein crystallization and crystallographic data collection

The PDLP5 ectodomain formed crystals in hanging drops composed of 1 μl of protein solution (70 mg/ml in 20 mM citrate pH 5.0 and 150 mM NaCl) and 1 μl of crystallization buffer (17.5 % [w/v] polyethylene glycol 4,000, 250 mM (NH_4_)_2_SO_4_) suspended over 800 μl of the latter as reservoir solution. Protein crystals were transferred into crystallization buffer supplemented with 25% (v/v) ethylene glycol, which served as cryoprotectant, and snap frozen in liquid N_2_. PDLP8 crystals (52 mg/ml in 20 mM citrate pH 5.0, 150 mM NaCl) developed in hanging drops containing 17.5 % (w/v) polyethylene glycol 4,000, 0.1 M citrate pH 5.5, 20% (v/v) 2-propanol. Crystals were frozen directly in liquid N_2_. For PDLP5 native (λ= 1.0 Å) and redundant sulfur SAD (λ= 2.079 Å) data were collected to 1.29 Å resolution at beam line PX-III of the Swiss Light Source (SLS), Villigen, Switzerland. A 1.95 Å native data set of PDLP8 was acquired at the same beam line. Data processing and reduction was done with XDS (version: Jan, 2018)^96^.

### Structure solution and refinement

The structure of PDLP5 was solved using the single-anomalous diffraction (SAD) method. 24 S sites corresponding to the 12 disulfide bonds in the PDLP5 crystallographic dimer were located with the program SHELXD^97^, site-refinement and phasing was done in SHARP^98^ and the starting phases were used for automated model building in BUCCANEER^99^ and ARP/wARP^100^. The model was completed in alternating cycles of model correction in COOT^101^ and restrained refinement in Refmac5^102^. The structure of PDLP8 was solved using the molecular replacement methods as implemented in the program PHASER^103^, and using the refined PDLP5 tandem ectodomain as search model. Inspection with MolProbity^104^ revealed excellent stereochemistry for the final models. Structural and surface representations were done in Pymol (http://pymol.soureforge.org) and Chimera^105^.

### Data availability

Materials used in this study and data generated are available from the corresponding author upon request. Phylogenetic trees with bootstrap information for 1000 replicates and corresponding sequence alignments have been deposited on Wasabi (http://wasabiapp.org); identifiers are available in the figure legends as web links. Information on used genomic data is available in Table S5. Publically available gene expression data was taken from the Sequence Read Archive (SRA) database; identifiers are listed in Table S4. Crystallographic coordinates and structure factors have been deposited with the Protein Data Bank (http://rcsb.org) with accession codes 6GRE (PDLP5) and 6GRF (PDLP8).

### Code availability

All R scripts developed for parsing the data and visualizing the results are available upon request.

## Supporting information

## Authors’ contributions

AV, BB, JK, JS, MH and MW conceived and designed the project. AL, BB, OS, SR, ML, AVe, AL, MH, and JS carried out the analyses. AV, BB, AL, MH, JS, and MW analyzed the data. AV, BB, MH, JS and MW wrote the manuscript. All authors read and contributed to the final manuscript.

## Acknowledgments

The authors would like to thank Ms. Kerri Hunter, Drs. Julia Krasensky-Wrzaczek, Sachie Kimura and Alexey Shapiguzov for critical comments on the manuscript. We thank Prof. David Collinge and Prof. Michael Lyngkjær (University of Copenhagen, Denmark) for help with the barley genome and Prof. Eric Schranz (Wageningen University, Netherlands) for helpful comments. This work was supported by the Doctoral Programme in Plant Sciences (DPPS) of the University of Helsinki and by the Finnish Cultural Foundation Suomen Kulttuurirahasto (to AV), the Academy of Finland (grant numbers #275632, #283139 and #312498 to MW) and the University of Helsinki (Three-year fund allocation to MW) and by grant 31003A_176237 from the Swiss National Science Foundation and by an International Research Scholar Award from the Howard Hughes Medical Institute (to MH). AV, OS, SR, JK, JS and MW are members of the Centre of Excellence in the Molecular Biology of Primary Producers (2014-2019) funded by the Academy of Finland (grant numbers #271832 and #307335). BB was supported by an EMBO long-term fellowship. We would also like to thank the staff at beam line PXIII of the Swiss Light Source (Villigen, Switzerland) for technical assistance during data collection.

---

**Figure S1. Summary of manual gene annotation and correction. a)** The number of corrected, manually annotated and partial/pseudo gene models in the studied species. Percentage of corrected gene models is marked with light gray, manually annotated genes with black and genes classified as partial or pseudogenes with dark gray. Silver birch (*Betula pendula*) and sacred lotus (*Nelumbo nucifera*) genes were fully manually annotated, as the gene models were not available when the study was initiated. *Selaginella moellendorffii* and *Vitis vinifera* required highest percentage of manual corrections. The high percentage of pseudogenes in *Physcomitrella patens* is explained by low gene number (two out of three gene models are likely pseudogenes). **b)** Average exon numbers of CRRSPs, PDLPs and CRKs. Average exon numbers were calculated for sdCRRSPs, ddCRRSPs, PDLPs and CRKs in *Amborella trichopoda*, *Arabidopsis thaliana* and *Oryza sativa*. **c)** The amount of curated and manually annotated gene models in basal and variable groups. Corrected (red) and manually annotated (green: species with pre-existing annotations; blue: species without previous annotations) gene models marked in both groups. Corrected or annotated genes can be found in all subgroups within these groups. There are several examples of corrected or previously non-annotated genes that are basal for subgroups, indicating the importance of gene model validation for correct tree topology.

**Figure S2. Phylogenies of DUF26-containing proteins. a)** A phylogenetic maximum-likelihood tree was estimated with full-length sequences for the basal group containing Selaginella sdCRRSPs and CRKs, Norway spruce CRRSPs and CRKs, monocot and eudicot bCRKs and PDLPs. Detailed phylogenetic trees with bootstrap support (1000 replicates) and filtered sequence alignment can be found at http://was.bi?id=wpEHGt. **b)** The phylogenetic maximum-likelihood tree for the variable group contains angiosperm CRRSPs and vCRKs. Tree was estimated using the full-length sequences. Detailed phylogenetic trees with bootstrap support (1000 replicates) and filtered sequence alignment can be found at http://was.bi?id=aIJe_D. Phylogenetic maximum likelihood trees of **c)** CRRSPs **d)** CRKs and **e)** PDLPs. Detailed phylogenetic trees containing gene identifiers as well as bootstrap support (1000 replicates) and filtered sequence alignment can be found at http://was.bi?id=zbIl7i (CRRSPs), http://was.bi?id=i9To8q (CRKs) and http://was.bi?id=Fe1A3A (PDLPs). **f)** Phylogenetic maximum-likelihood tree of all DUF26 genes in *Marchantia polymorpha*, *Selaginella moellendorffii* and *Amborella trichopoda*. Tree is estimated from sequence alignment of full length gene models where the sites with coverage less than 10% have been filtered out. Tree is rooted to sdCRRSPs from *Marchantia polymorpha*. A detailed phylogenetic tree with gene identifiers as well as bootstrap support (1000 replicates) and filtered sequence alignment can be found at http://was.bi?id=VeeQZ6.

**Figure S3. Ancestral gene counts for DUF26-containing genes.** DLCpar was used for inferring the most parsimonious history of protein groups in the presence of duplications, losses, and incomplete lineage sorting. The panels illustrate ancestral gene counts and lineage-specific expansions in **a)** sdCRRSPs in the basal group, **b)** basal CRKs, **c)** PDLPs, **d)** variable group CRKs, and **e)** ddCRRSPs in the variable group. Numbers with black color show the gene counts in the species and their most recent common ancestor. Estimated gene gains are marked with red and losses with blue.

**Figure S4. Badirate comparisons for evolutionary rates.** Analyses were carried out with Badirate for eleven species (*Physcomitrella patens*, *Selaginella moellendorffii*, *Amborella trichopoda*, *Arabidopsis thaliana*, *Populus trichocarpa*, *Vitis vinifera*, *Aquilegia coerulea*, *Spirodela polyrhiza*, *Zea mays*, *Oryza sativa* and *Brachypodium distachyon*). Neutral branches: bold black lines; gene family expansion: bold purple lines; gene family contraction: blue dashed lines. Branches with a significant difference to birth-death model estimated from orthogroup data are marked with arrows. Node labels present the gene family size in ancestral nodes as estimated by Badirate. Tip labels contain species abbreviation and the change in number compared to the most recent ancestral node. **a)** All CRKs compared to all kinases. **b)** bCRKs compared to all kinases. **c)** vCRKs compared to all kinases.

**Figure S5. Phylogenetic maximum-likelihood tree of bCRKs.** The full length sequences belonging to this clade were re-aligned and the alignment was filtered to exclude sites with less than 10% coverage. Bootstrap support is calculated with 1000 replicates. A detailed phylogenetic tree and filtered sequence alignment can be found at http://was.bi?id=6Z7yhQ.

**Figure S6. Species trees and reconciled phylogenetic trees for DCLpar analyses. a)** Species tree for the 24 species where all DUF26-domain genes were comprehensively analyzed. The tree was downloaded from PhyloT. The node labels indicate the speciation event IDs that are used in panels b and c. **b)** Reconciled gene tree for the bCRKs from DCLpar. The node labels provide the timing of the event by referring to the speciation event ID in the species tree. **c)** Reconciled gene tree for the variable group CRRSPs from DLCpar. The node labels provide the timing of the event by referring to the speciation event ID in the species tree.

**Figure S7. Phylogenetic maximum-likelihood tree of 5 species used in segmental duplication analyses and *Selaginella moellendorffii* as outgroup.** The tree includes DUF26 genes from *Amborella trichopoda*, *Solanum lycopersicum*, *Arabidopsis thaliana*, *Oryza sativa*, *Zea mays* and *Selaginella moellendorffii*. The full length gene models were used for the sequence alignment and the sites with less than 10% coverage were filtered out. Bootstrap support is calculated with 1000 replicates. A detailed phylogenetic tree and filtered sequence alignment can be found at http://was.bi?id=2NeJCb.

**Figure S8. Phylogenetic maximum-likelihood tree of PDLPs with possible partial PDLP from *Marsilea quadrifolia*.** The phylogenetic tree is based on the sequence covering the part of ectodomain that is present in the partial gene model from *Marsilea quandrifolia*. The ddCRKs from *Selaginella moellendorffii*, *Picea abies* and *Amborella trichopoda* were used as outgroup for PDLPs. The partial gene model from fern *Marsilea quadrifolia* is placed close to the root of PDLP clade and thus could be a PDLP. Bootstrap support is calculated with 1000 replicates. A detailed phylogenetic tree and filtered sequence alignment can be found at http://was.bi?id=usJEbx.

**Figure S9: Mutation of disulfide bridge-forming cysteines in PDLP5 results in protein aggregation.** PDLP5, PDLP5^C101A^, PDLP5^C148A^ and PDLP5^C191A^ ectodomains were subjected to preparative size exclusion chromatography (left). Non-aggregated fractions were combined and subjected to analytical size exclusion chromatography (right). Molecular mass standards: A = Thyroglobulin, 669 kDa B = Aldolase, 158 kDa; C = Conalbumin, 75 kDa; D = Ovalbumin, 44 kDa; E = Ribonuclease A, 13.7 kDa.

**Figure S10: Mutations in disulfide bridge forming residues in PDLP5 result in lower protein stability:** Melting curves (4 replicates in green, brown, red and blue) of PDLP5, PDLP5^C101A^, PDLP5^C148A^, PDLP5^C191A^ ectodomains and of the blank without protein (blank measurements for PDLP5, PDLP5^C101A^, PDLP5^C148A^ are the same as the experiments were carried out together). For PDLP5, PDLP5^C101A^, PDLP5^C148A^ ectodomains average melting temperatures are given +/- SDM (n=4). PDLP5^C191A^ was unstable at the given conditions and no melting curve could be acquired.

**Figure S11: Structural comparisons of PDLP5 and PDLP8 DUF26 domains reveal a high degree of structural similarity (a)** Superimposition of the DUF26-A (orange; C_α_ trace) and the DUF26-B (blue; C_α_ trace) domains of PDLP5 (left; r.m.s.d. is ~1.6 Å comparing 89 corresponding C_α_ atoms) and PDLP8 (right; r.m.s.d. is ~1.2 Å comparing 89 corresponding C_α_ atoms) demonstrate the structural similarity of DUF26-A and DUF26-B domains. Glycosylated asparagines are indicated by an arrow **(b)** Structural superposition of PDLP5 (orange, shown as C_α_ trace) and PDLP8 (blue) reveals a high degree of overall structural similarity (r.m.s.d. is ~1.6 Å comparing 198 corresponding C_α_ atoms), and a conserved pattern of disulfide bridges (grey highlights). The disulfide bridges in PDLP8 are: 1 (Cys89-Cys98), 2 (Cys101-Cys126), 3 (Cys34-Cys113), 4 (Cys191-Cys200), 5 (Cys203-Cys228) and 6 (Cys148-Cys215). Disulfide bridges are depicted in bonds representation (PDLP5 in yellow, PDLP8 in green).

**Figure S12: Cysteines forming disulfide bonds and residues involved in the interaction of DUF26-A and DUF26-B domains are conserved in bCRKs, vCRKs, CRSPs and PDLPs.** A set of PDLPs, bCRKs, vCRKs and CRRSPs were selected based on the structure and their sequences were aligned with MUSCLE^106^. The result shows the conservation of amino acids present in the interaction patch of DUF26-A and DUF26-B in either PDLP5s (red highlight) or all double DUF26 containing proteins (highlight in blue). Cysteines and disulfide bridges are highlighted in yellow. A secondary structure assignment of the DUF26-A (blue) and DUF26-B domains^107^ is given above the sequences.

**Figure S13: The PDLP5 ectodomain does not bind mannose or other cell wall derived sugars. a)** Mannose was titrated into a cell containing the PDLP5 ectodomain in an isothermal titration calorimetry (ITC) assay (n.d., no binding detected). **b)** ITC experiments were carried out to test binding of plant cell wall sugars to the isolated PDLP5 ectodomain.

**Figure S14. Gene tree reconciliation of the five species used in segmental duplication analyses. (a)** Species tree of the five species in the analyses. The phylogeny was downloaded from PhyloT, and the node labels indicate the speciation event ID. These IDs are used in Figures S15b-d). The reconciled gene trees were estimated with DLCpar for **(b)** bCRKs, **(c)** vCRKs, **(d)** ddCRRSPs. The node labels provide the timing of the split by referring to the speciation event ID in the species tree (Figure S15a). *Selaginella moellendorffii* was used as outgroup.

**Figure S15. The DUF26 genes show transcriptional response to several stress treatments.** Heatmap illustrating transcriptional response of DUF26 genes from *Arabidopsis thaliana*, *Oryza sativa*, *Zea mays* and *Solanum lycopersicum*. The dendrogram shows a phylogenetic tree of the 253 DUF26-containing genes (rows) in the four species. The columns represent the RNAseq experiments from Sequence Read Archive (see Table S4; accession numbers not shown here for clarity), categorized into pathogen defence (red highlight) and miscellaneous (blue). The heatmap colors represent the log**2**(TPM) values, as illustrated by the color key. The NA values are displayed with white color.

**Supplementary file 1: Sequences used for analyses in fasta format**

**Supplementary file 2: wwPDB X-ray Structure Validation Summary Report 6GRE (PDLP5)**

**Supplementary file 3: wwPDB X-ray Structure Validation Summary Report 6GRF (PDLP8)**

**Table S1: Information of DUF26 proteins included in this study.** Information of DUF26 protein sequences found in study species.

**Table S2: Data collection, phasing and refinement statistics for structural analyses**

**Table S3: The related kinase domains for the CRK kinase domains.** PBLAST results for selected CRKs from *Amborella trichopoda*, *Arabidopsis thaliana*, *Oryza sativa* and *Selaginella moellendorffii*. Only amino acid sequence of the kinase domain of each CRK was used as query. Best hit outside the CRKs was marked in the table.

**Table S4: Gene conversion analyses results.**

**Table S5: Identified orthologs and information of transcriptome data used in analyses.**

**Table S6: Genome version information and references for plant genomes used in phylogenetic analyses.**

