## Supplementary material for "Mechanistic insights into the evolution of DUF26-containing proteins in land plants"

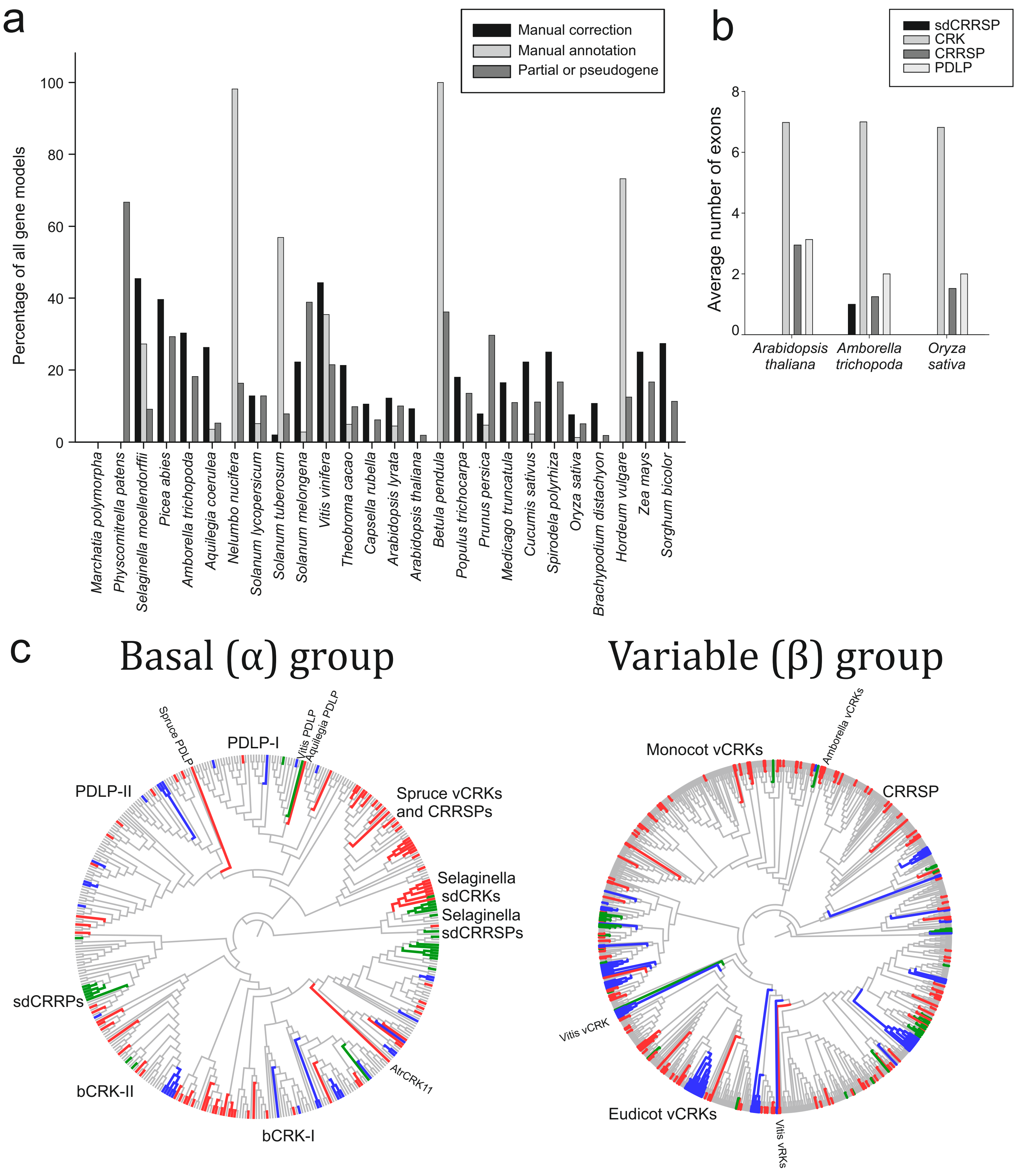


**Figure S1. Summary of manual gene annotation and correction. a)** The number of corrected, manually annotated and partial/pseudo gene models in the studied species. Percentage of corrected gene models is marked with light grey, manually annotated genes with black and genes classified as partial or pseudogenes with dark grey. Silver birch (*Betula pendula*) and sacred lotus (*Nelumbo nucifera*) genes were fully manually annotated, as the gene models were not available when the study was initiated. *Selaginella moellendorffii* and *Vitis vinifera* required highest percentage of manual corrections. The high percentage of pseudogenes in *Physcomitrella patens* is explained by low gene number (two out of three gene models are likely pseudogenes). **b)** Average exon numbers of CRRSPs, PDLPs and CRKs. Average exon numbers were calculated for sdCRRSPs, ddCRRSPs, PDLPs and CRKs in *Amborella trichopoda*, *Arabidopsis thaliana* and *Oryza sativa*. **c)** The amount of curated and manually annotated gene models in basal and variable groups. Corrected (red) and manually annotated (green: species with pre-existing annotations; blue: species without previous annotations) gene models marked in both groups. Corrected or annotated genes can be found in all subgroups within these groups. There are several examples of corrected or previously non-annotated genes that are basal for subgroups, indicating the importance of gene model validation for correct tree topology.

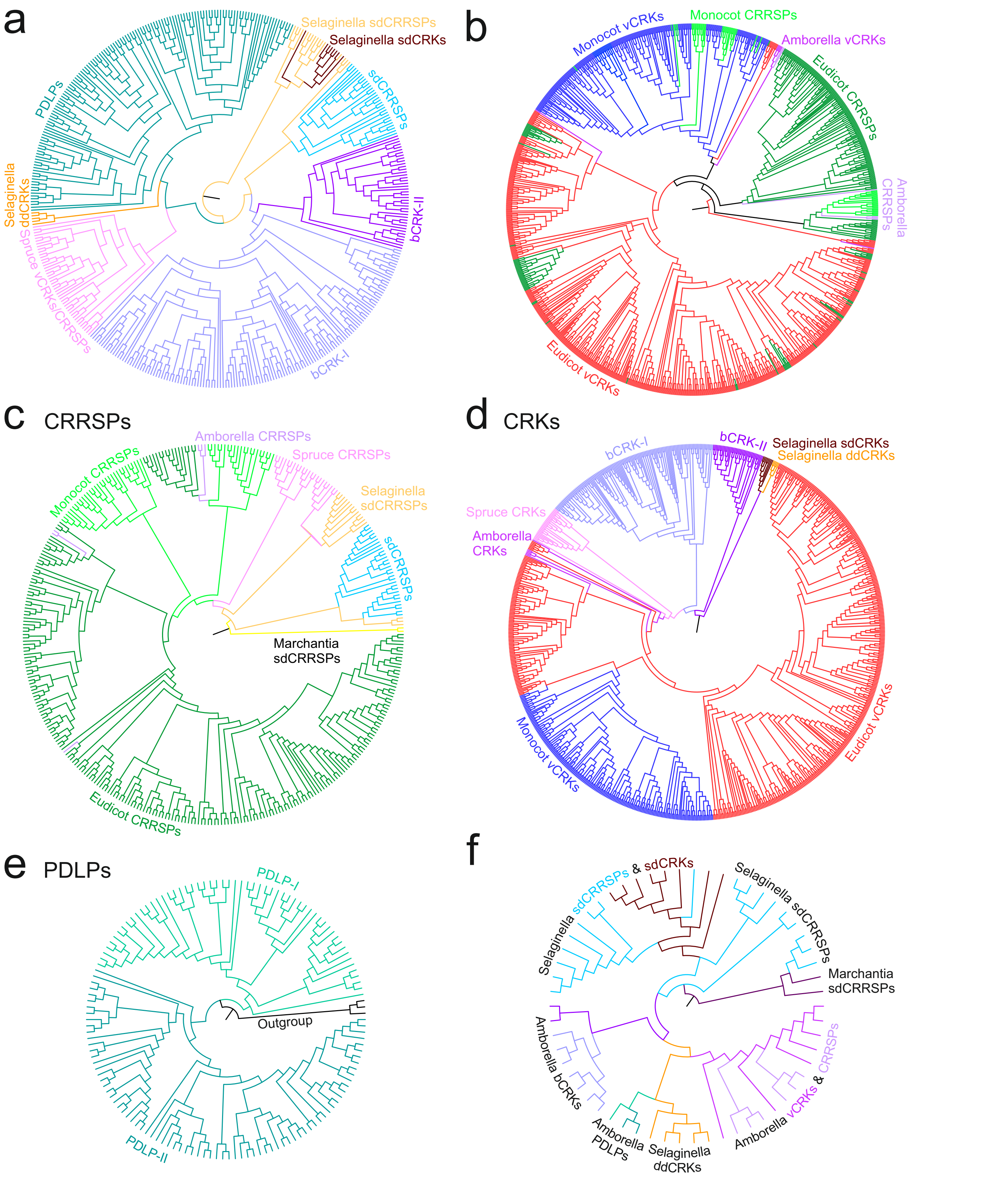


**Figure S2. Phylogenies of DUF26-containing proteins. a)** A phylogenetic maximum-likelihood tree was estimated with full-length sequences for the basal group containing Selaginella sdCRRSPs and CRKs, Norway spruce CRRSPs and CRKs, monocot and eudicot bCRKs and PDLPs. Detailed phylogenetic trees with bootstrap support (1000 replicates) and filtered sequence alignment can be found at http://was.bi?id=wpEHGt. **b)** The phylogenetic maximum-likelihood tree for the variable group contains angiosperm CRRSPs and vCRKs. Tree was estimated using the full-length sequences. Detailed phylogenetic trees with bootstrap support (1000 replicates) and filtered sequence alignment can be found at http://was.bi?id=aIJe_D. Phylogenetic maximum likelihood trees of **c)** CRRSPs **d)** CRKs and **e)** PDLPs. Detailed phylogenetic trees containing gene identifiers as well as bootstrap support (1000 replicates) and filtered sequence alignment can be found at http://was.bi?id=zbIl7i (CRRSPs), http://was.bi?id=i9To8q (CRKs) and http://was.bi?id=Fe1A3A (PDLPs). **f)** Phylogenetic maximum-likelihood tree of all DUF26 genes in *Marchantia polymorpha*, *Selaginella moellendorffii* and *Amborella trichopoda*. Tree is estimated from sequence alignment of full length gene models where the sites with coverage less than 10% have been filtered out. Tree is rooted to sdCRRSPs from *Marchantia polymorpha*. A detailed phylogenetic tree with gene identifiers as well as bootstrap support (1000 replicates) and filtered sequence alignment can be found at http://was.bi?id=VeeQZ6.

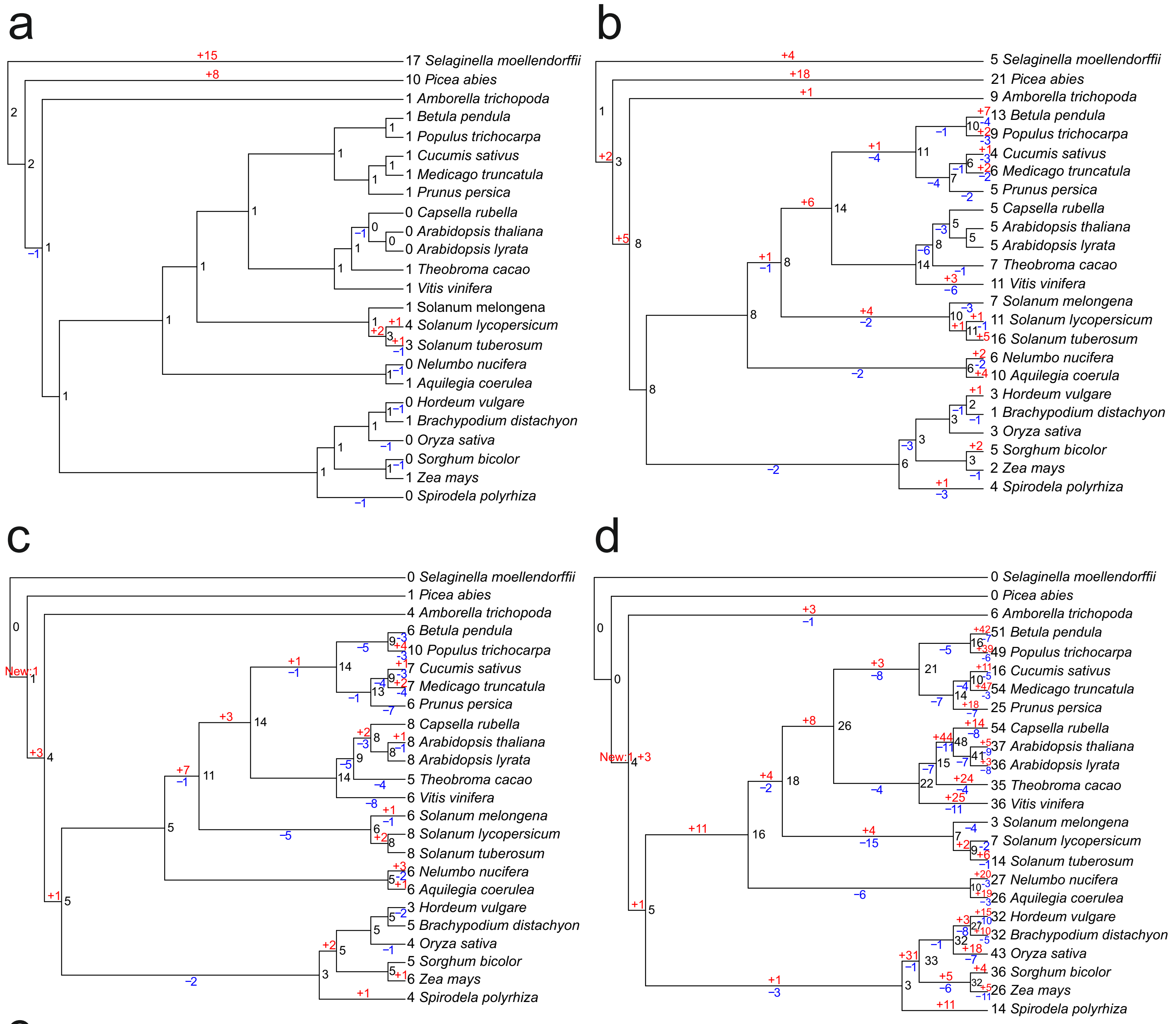


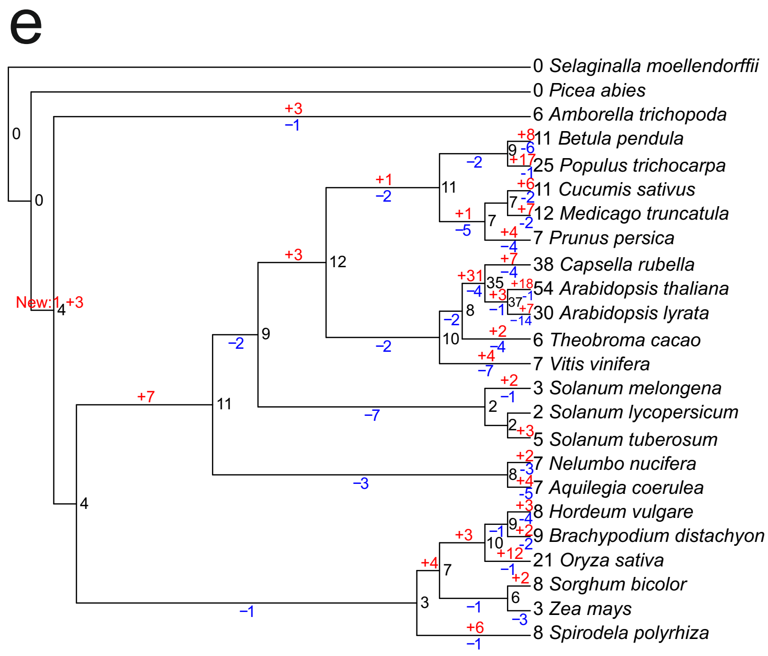


**Figure S3. Ancestral gene counts for DUF26-containing genes.** DLCpar was used for inferring the most parsimonious history of protein groups in the presence of duplications, losses, and incomplete lineage sorting. The panels illustrate ancestral gene counts and lineage-specific expansions in **a)** sdCRRSPs in the basal group, **b)** basal CRKs, **c)** PDLPs, **d)** variable group CRKs and **e)** ddCRRSPs in the variable group. Numbers with black color show the gene counts in the species and their most recent common ancestor. Estimated gene gains are marked with red and losses with blue.

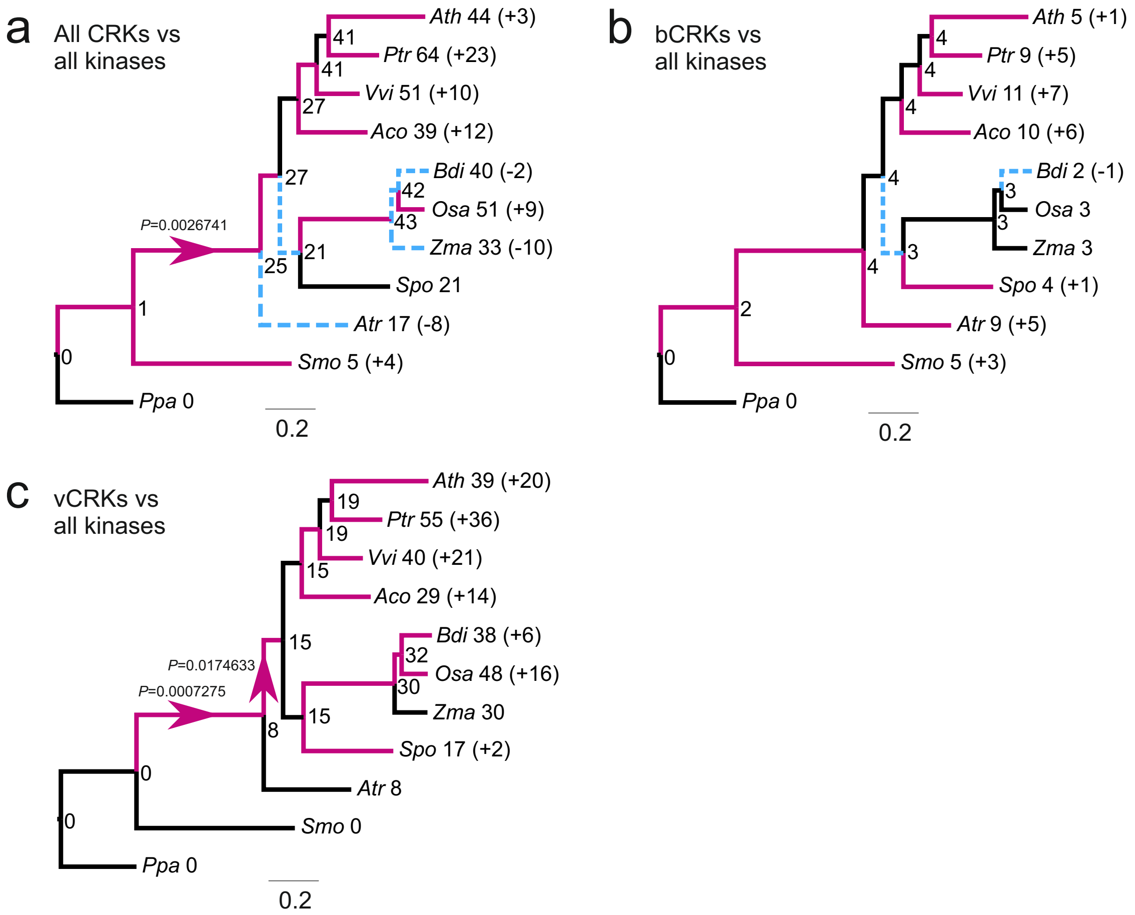


**Figure S4. Badirate comparisons for evolutionary rates.** Analyses were carried out with Badirate for eleven species (*Physcomitrella patens*, *Selaginella moellendorffii*, *Amborella trichopoda*, *Arabidopsis thaliana*, *Populus trichocarpa*, *Vitis vinifera*, *Aquilegia coerulea*, *Spirodela polyrhiza*, *Zea mays*, *Oryza sativa* and *Brachypodium distachyon*). Neutral branches: bold black lines; gene family expansion: bold purple lines; gene family contraction: blue dashed lines. Branches with a significant difference to birth-death model estimated from orthogroup data are marked with arrows. Node labels present the gene family size in ancestral nodes as estimated by Badirate. Tip labels contain species abbreviation and the change in number compared to the most recent ancestral node. **a)** All CRKs compared to all kinases. **b)** bCRKs compared to all kinases. **c)** vCRKs compared to all kinases.

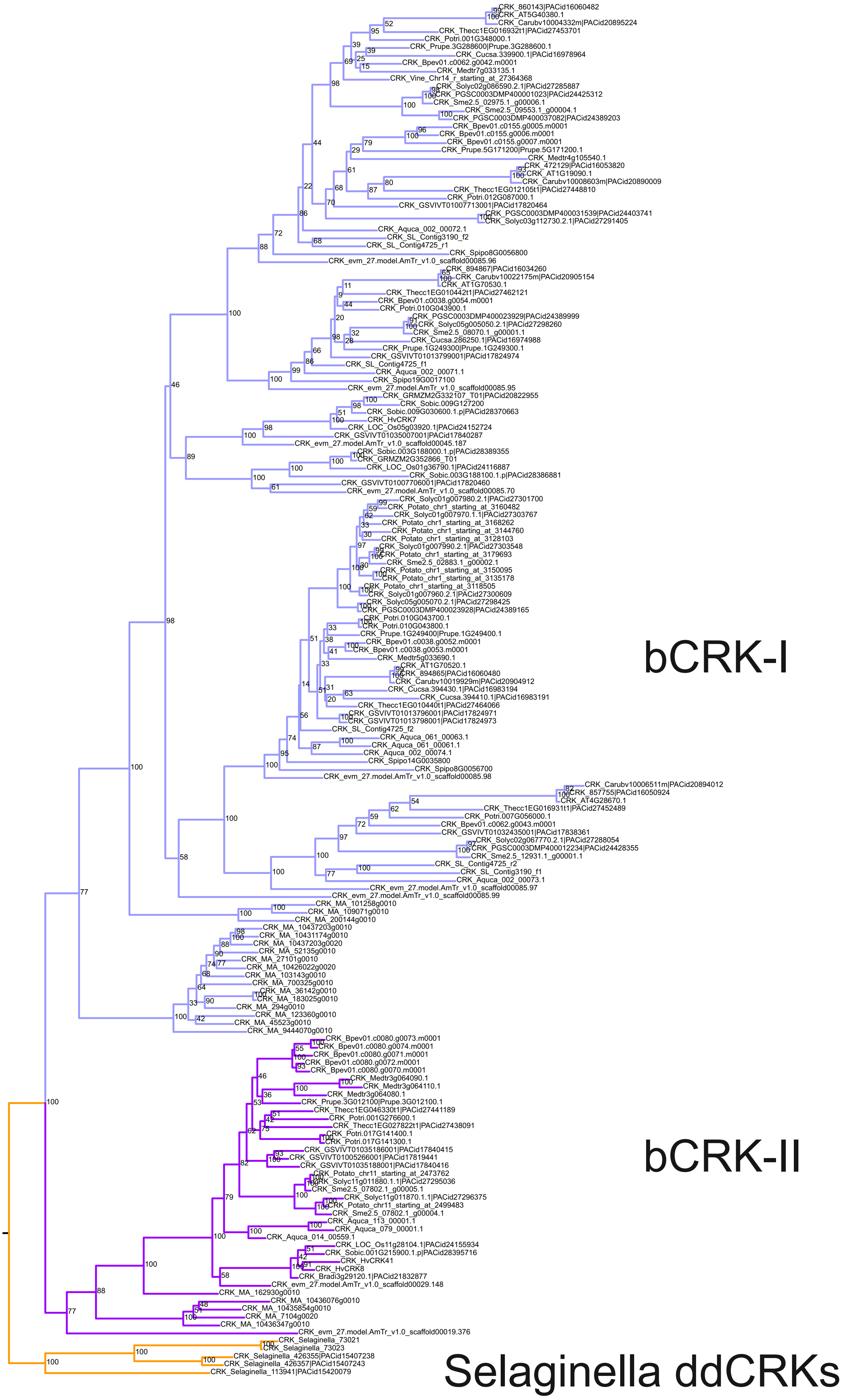


**Figure S5. Phylogenetic maximum-likelihood tree of bCRKs.** The full length sequences belonging to this clade were re-aligned and the alignment was filtered to exclude sites with less than 10% coverage. Bootstrap support is calculated with 1000 replicates. A detailed phylogenetic tree and filtered sequence alignment can be found at [http://was.bi?id=6Z7yhQ](http://was.bi?id=6Z7yhQ).

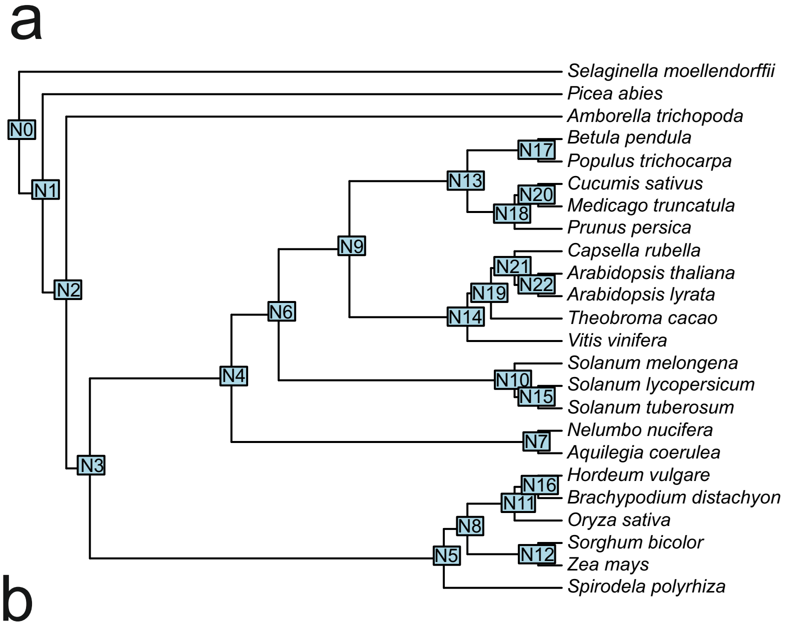


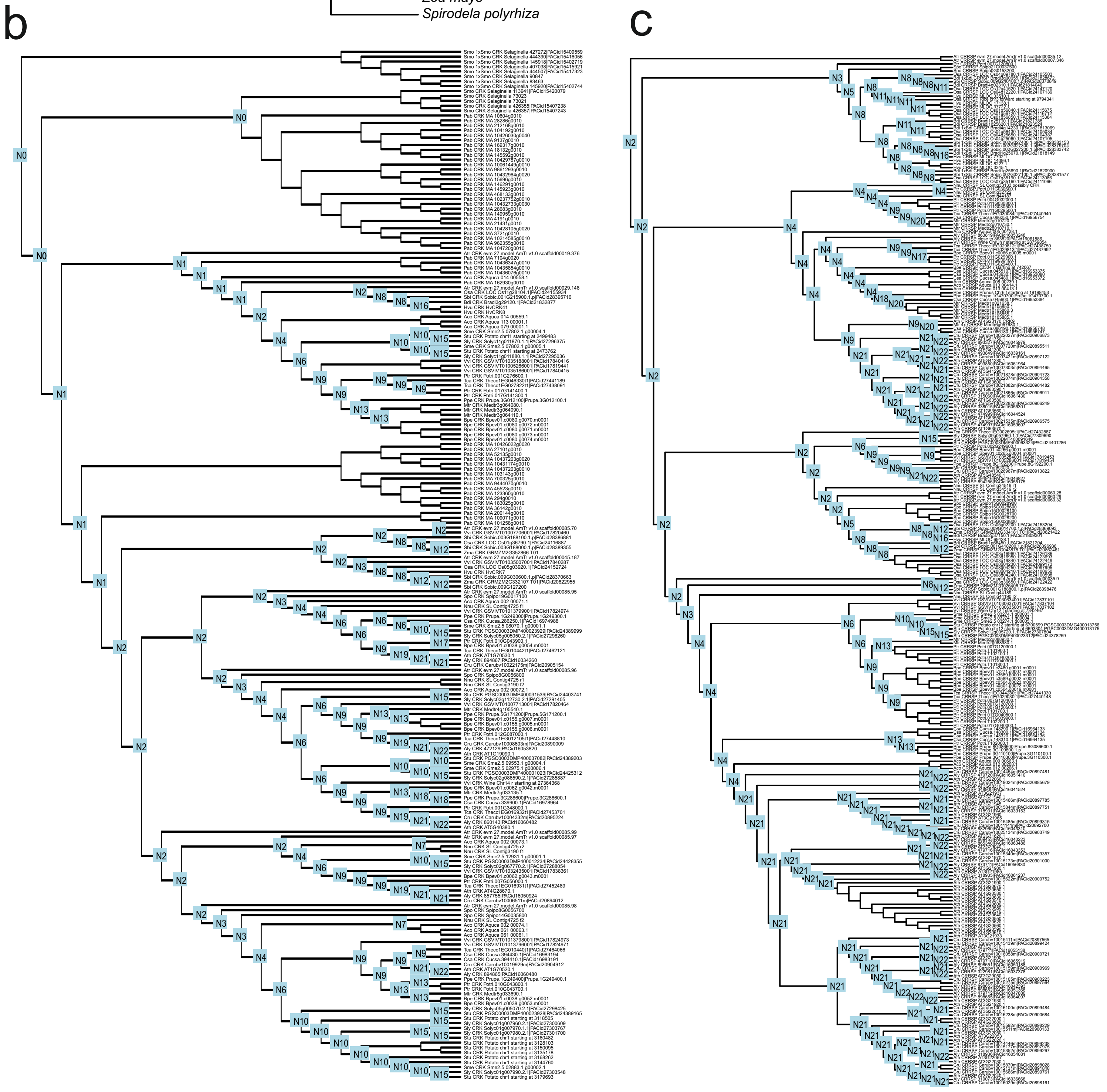


**Figure S6. Species trees and reconciled phylogenetic trees for DCLpar analyses. a)** Species tree for the 24 species where all DUF26-domain genes were comprehensively analyzed. The tree was downloaded from PhyloT. The node labels indicate the speciation event IDs that are used in panels b and c. **b)** Reconciled gene tree for the bCRKs from DCLpar. The node labels provide the timing of the event by referring to the speciation event ID in the species tree. **c)** Reconciled gene tree for the variable group CRRSPs from DLCpar. The node labels provide the timing of the event by referring to the speciation event ID in the species tree.

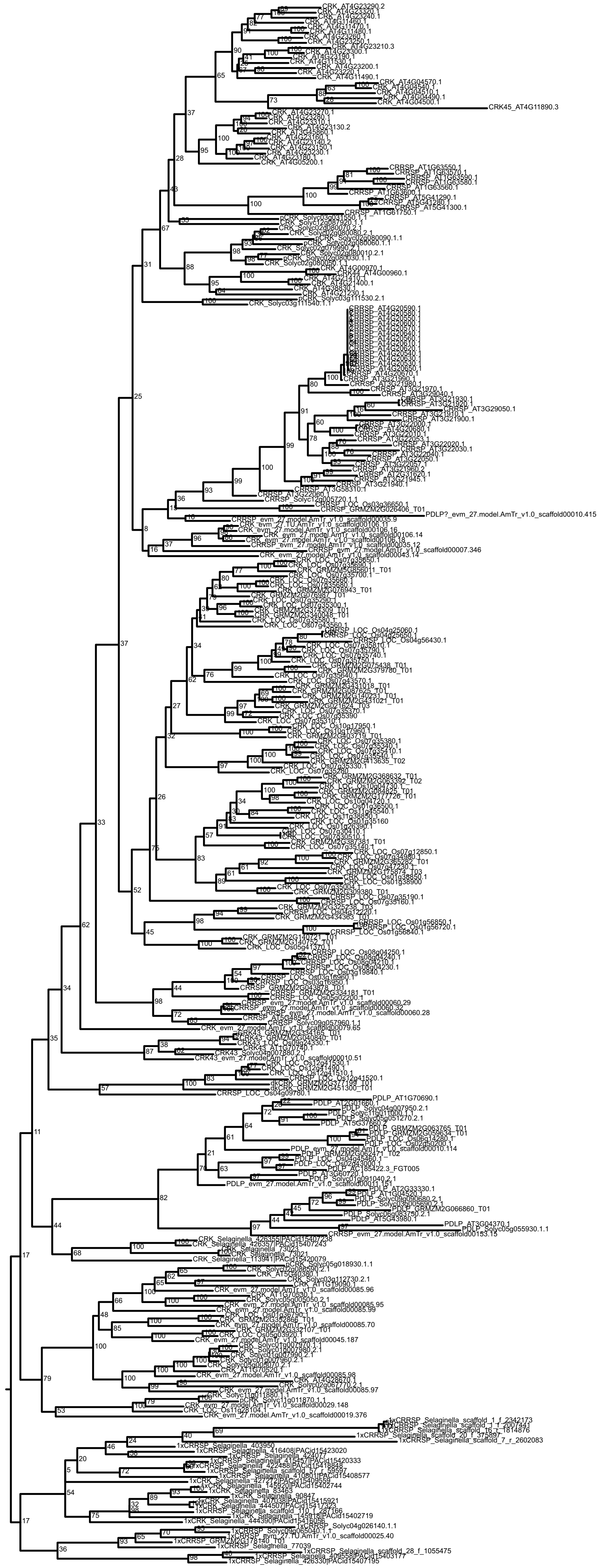

**Figure S7. Phylogenetic maximum-likelihood tree of 5 species used in segmental duplication analyses and *Selaginella moellendorffii* as outgroup.** The tree includes DUF26 genes from *Amborella trichopoda*, *Solanum lycopersicum*, *Arabidopsis thaliana*, *Oryza sativa*, *Zea mays* and *Selaginella moellendorffii*. The full length gene models were used for the sequence alignment and the sites with less than 10% coverage were filtered out. Bootstrap support is calculated with 1000 replicates. A detailed phylogenetic tree and filtered sequence alignment can be found at http://was.bi?id=2NeJCb.

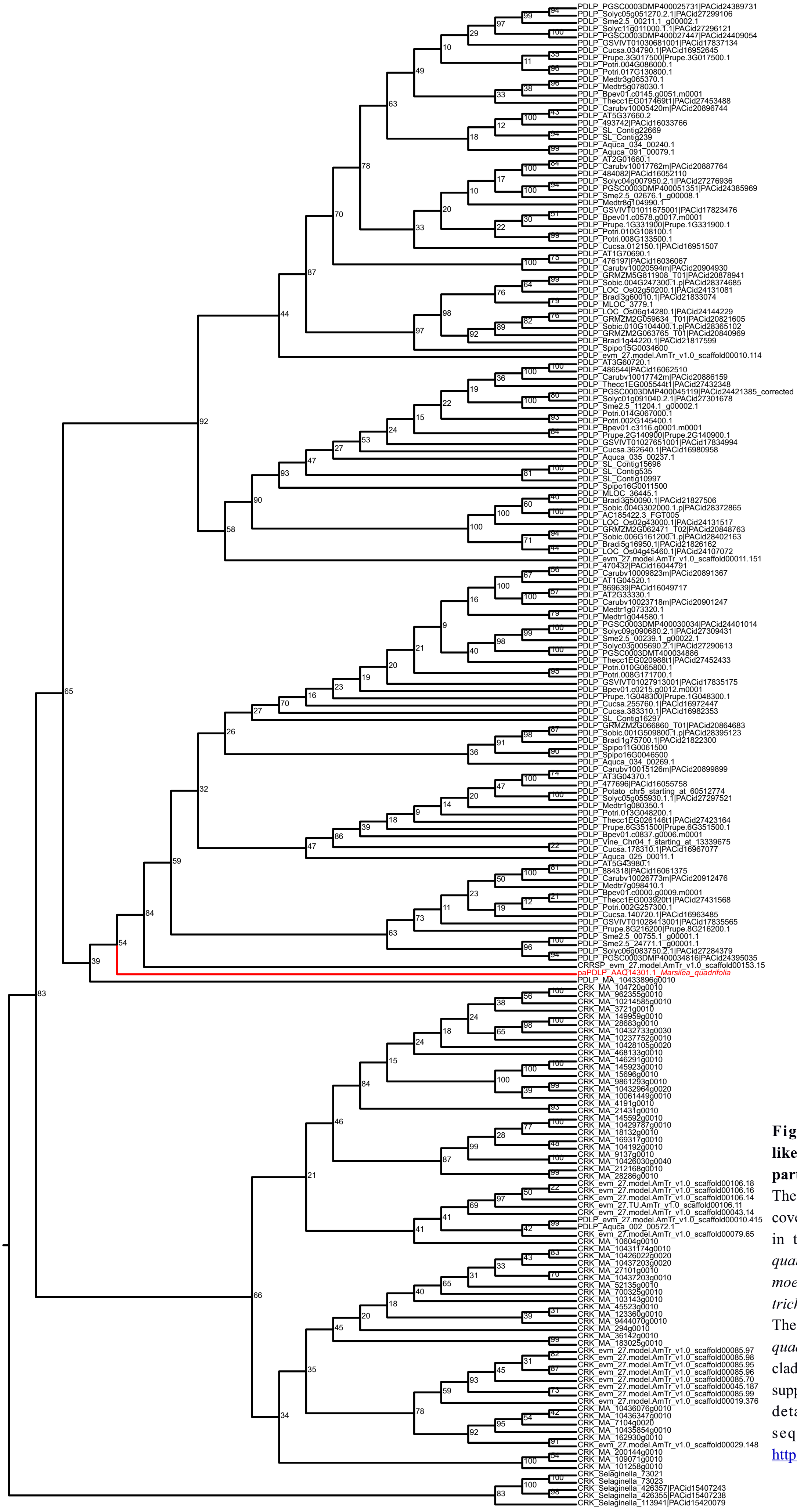


**Figure S8. Phylogenetic maximum-likelihood tree of PDLPs with possible partial PDLP from *Marsilea quadrifolia*.** The phylogenetic tree is based on the sequence covering the part of ectodomain that is present in the partial gene model from *Marsilea quandrifolia*. The ddCRKs from *Selaginella moellendorffii*, *Picea abies* and *Amborella trichopoda* were used as outgroup for PDLPs. The partial gene model from fern *Marsilea quadrifolia* is placed close to the root of PDLP clade and thus could be a PDLP. Bootstrap support is calculated with 1000 replicates. A detailed phylogenetic tree and filtered sequence alignment can be found at http://was.bi?id=usJEbx.

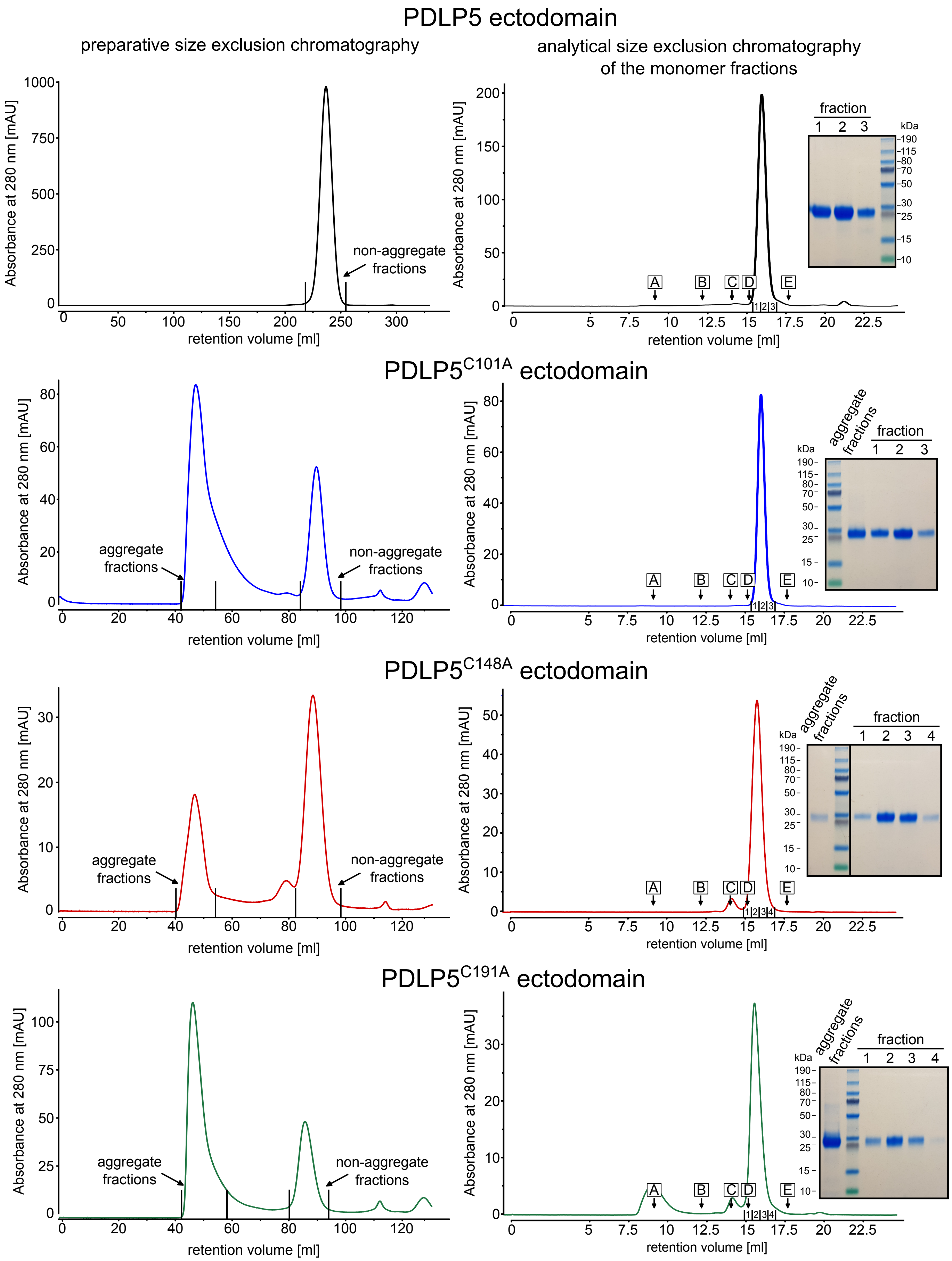


**Figure S9: Mutation of disulfide bridge forming cysteines in PDLP5 results in protein aggregation.** PDLP5, PDLP5$^{C101A}$, PDLP5$^{C148A}$ and PDLP5$^{C191A}$ ectodomains were subjected to preparative size exclusion chromatography (left). Non-aggregated fractions were combined and subjected to analytical size exclusion chromatography (right). Molecular mass standards: A = Thyroglobulin, 669 kDa B = Aldolase, 158 kDa; C = Conalbumin, 75 kDa; D = Ovalbumin, 44 kDa; E = Ribonuclease A, 13.7 kDa.

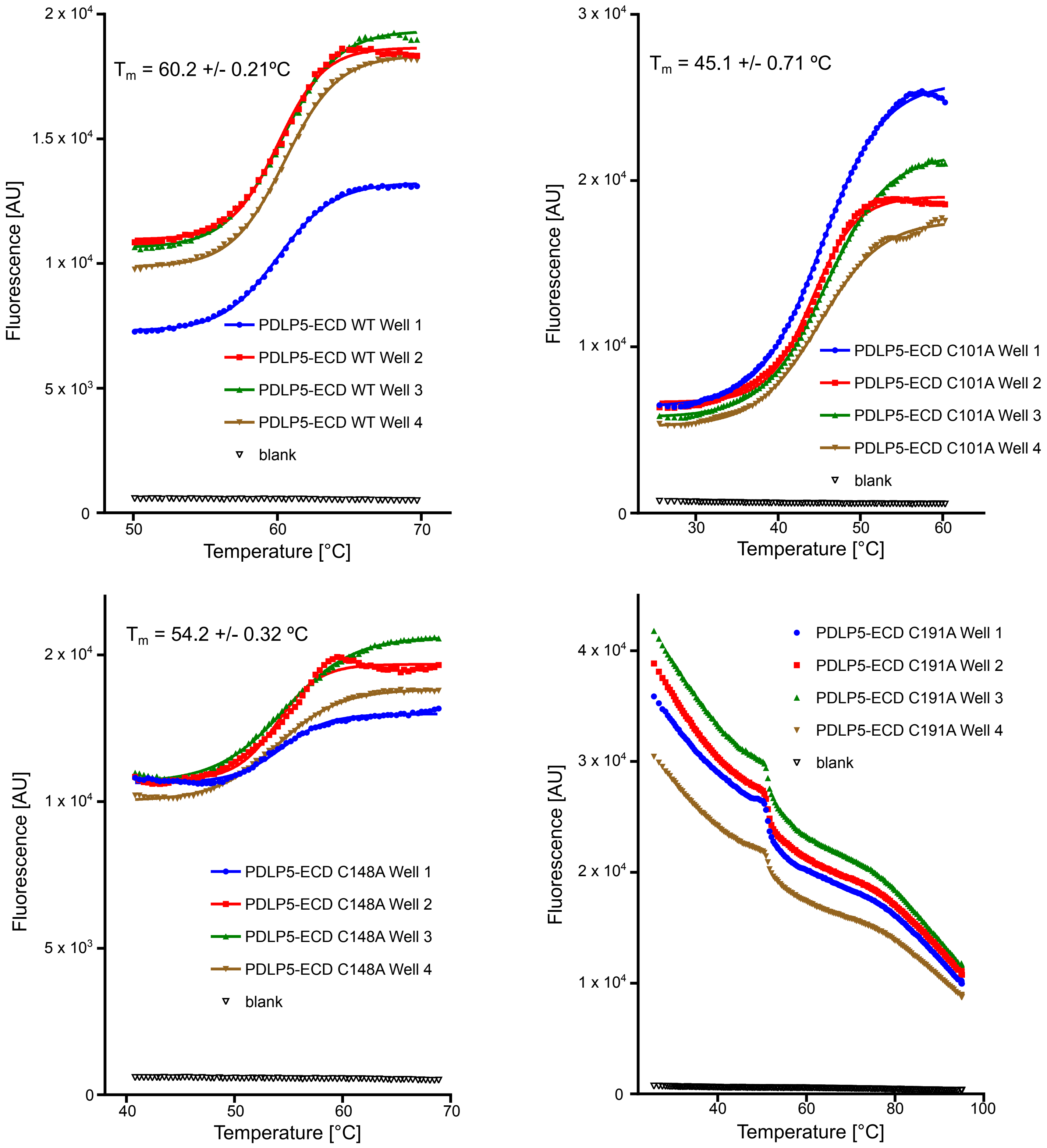


**Figure S10: Mutations in disulfide bridge forming residues in PDLP5 result in lower protein stability:** Melting curves (4 replicates in green, brown, red and blue) of PDLP5, PDLP5$^{C101A}$, PDLP5$^{C148A}$, PDLP5$^{C191A}$ ectodomains and of the blank without protein (blank measurements for PDLP5, PDLP5$^{C101A}$, PDLP5$^{C148A}$ are the same as the experiments were carried out together). For PDLP5, PDLP5$^{C101A}$, PDLP5$^{C148A}$ ectodomains average melting temperatures are given +/- SDM (n=4). PDLP5$^{C191A}$ was unstable at the given conditions and no melting curve could be acquired.

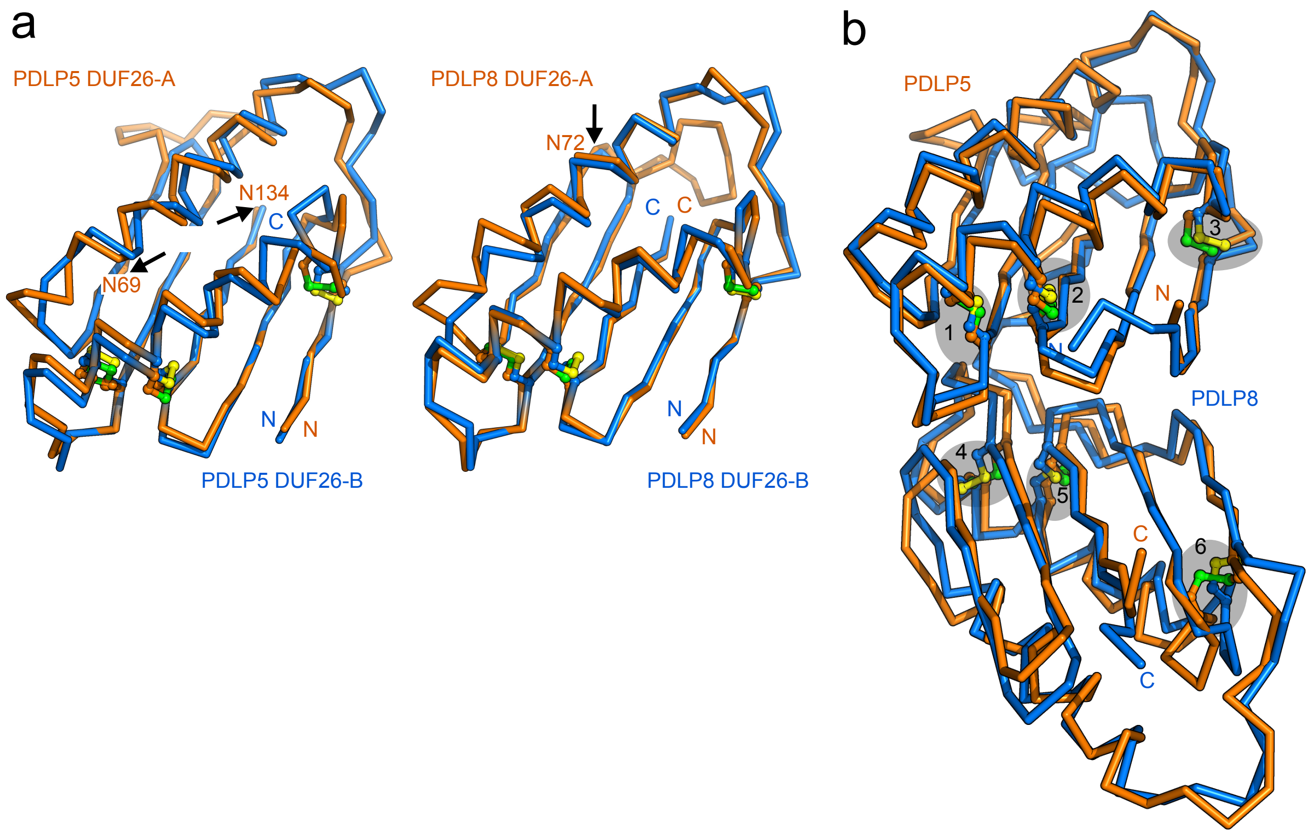


**Figure S11: Structural comparisons of PDLP5 and PDLP8 DUF26 domains reveal a high degree of structural similarity (a)** Superimposition of the DUF26-A (orange; $C_\alpha$ trace) and the DUF26-B (blue; $C_\alpha$ trace) domains of PDLP5 (left; r.m.s.d. is ~1.6 Å comparing 89 corresponding Cα atoms) and PDLP8 (right; r.m.s.d. is ~1.2 Å comparing 89 corresponding $C_\alpha$ atoms) demonstrate the structural similarity of DUF26-A and DUF26-B domains. Glycosylated asparagines are indicated by an arrow **(b)** Structural superposition of PDLP5 (orange, shown as $C_\alpha$ trace) and PDLP8 (blue) reveals a high degree of overall structural similarity (r.m.s.d. is ~1.6 Å comparing 198 corresponding $C_\alpha$ atoms), and a conserved pattern of disulfide bridges (grey highlights). The disulfide bridges in PDLP8 are: 1 (Cys89-Cys98), 2 (Cys101-Cys126), 3 (Cys34-Cys113), 4 (Cys191- Cys200), 5 (Cys203-Cys228) and 6 (Cys148-Cys215). Disulfide bridges are depicted in bonds representation (PDLP5 in yellow, PDLP8 in green).

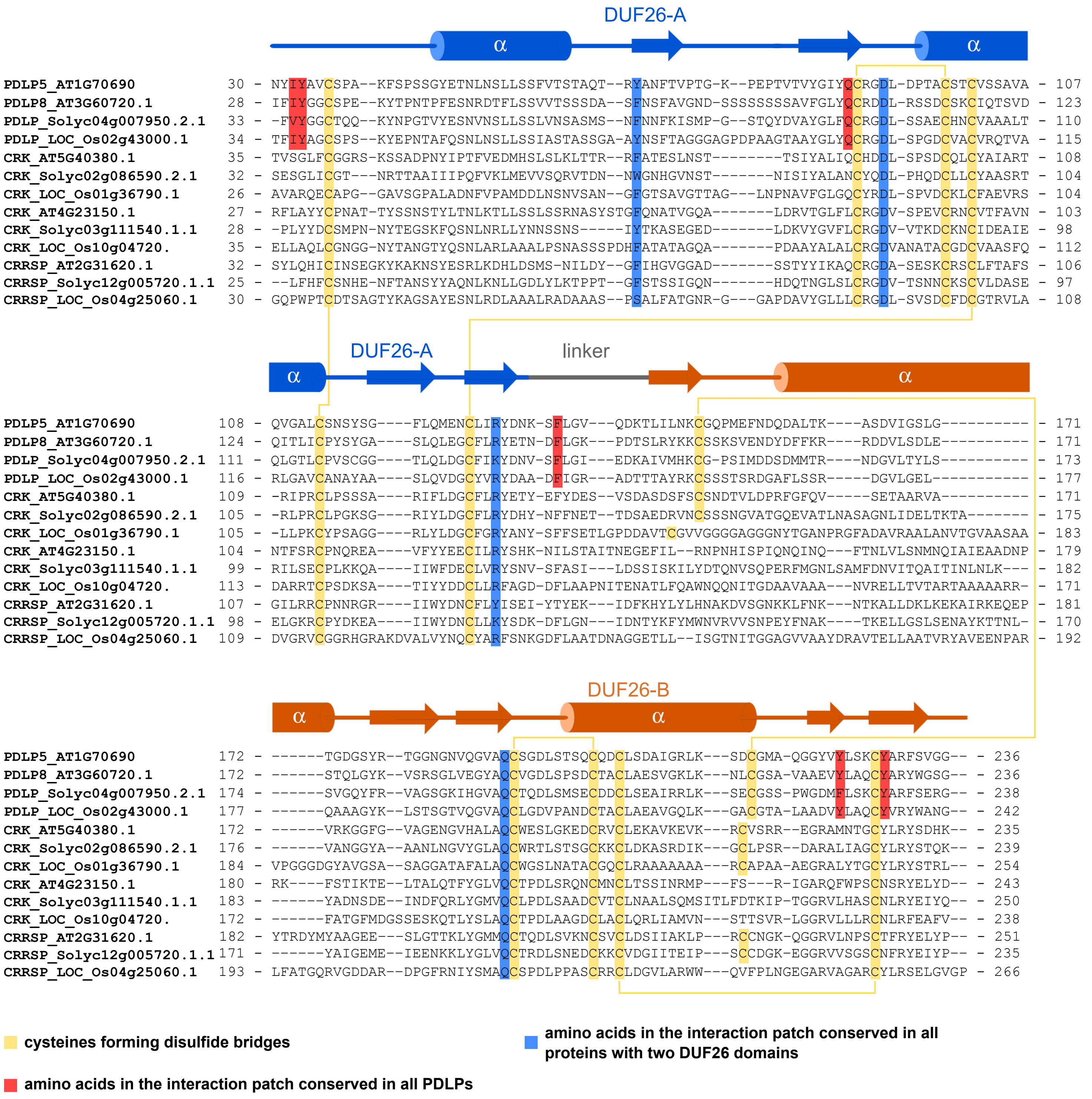


**Figure S12: Cysteines forming disulfide bonds and residues involved in the interaction of DUF26-A and DUF26-B domains are conserved in bCRKs, vCRKs, CRSPs and PDLPs.** A set of PDLPs, bCRKs, vCRKs and CRRSPs were selected based on the structure and their sequences were aligned with MUSCLE[90]. The result shows the conservation of amino acids present in the interaction patch of DUF26-A and DUF26-B in either PDLP5s (red highlight) or all double DUF26 containing proteins (highlight in blue). Cysteines and disulfide bridges are highlighted in yellow. A secondary structure assignment of the DUF26-A (blue) and DUF26-B domains[91] is given above the sequences.

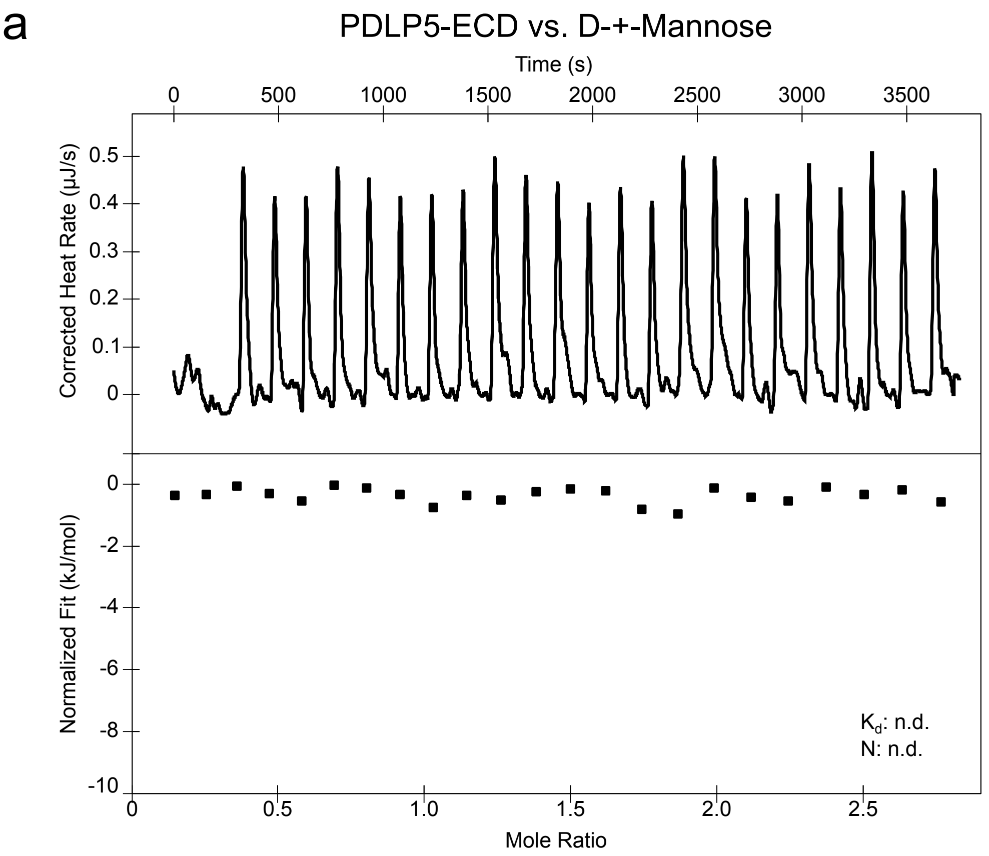


b

cell wall occuring sugars tested for binding to PDLP5-ECD by ITC

| | |
|---|---|
| Pectic Galactan | no binding detected |
| Rhamnogalacturonan | no binding detected |
| Polygalacturonic Acid | no binding detected |
| Cellohexaose | no binding detected |
| Arabinohexaose | no binding detected |

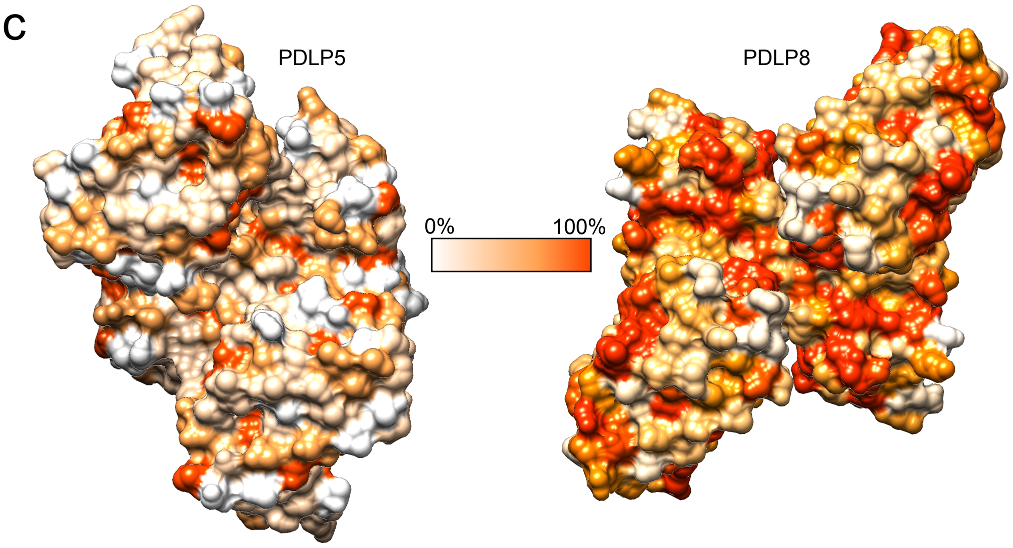


**Figure S13: The PDLP5 ectodomain does not bind mannose or other cell wall derived sugars and PDLP5 as well as PDLP8 surface exposed residues are not widely conserved. (a)** Mannose was titrated into a cell containing the PDLP5 ectodomain in an isothermal titration calorimetry (ITC) assay (n.d., no binding detected). **(b)** ITC experiments were carried out to test binding of plant cell wall sugars to the isolated PDLP5 ectodomain. **(c)**The conservation of amino acids is depicted on the surface of PDLP5 (left) or PDLP8 (left), respectively.

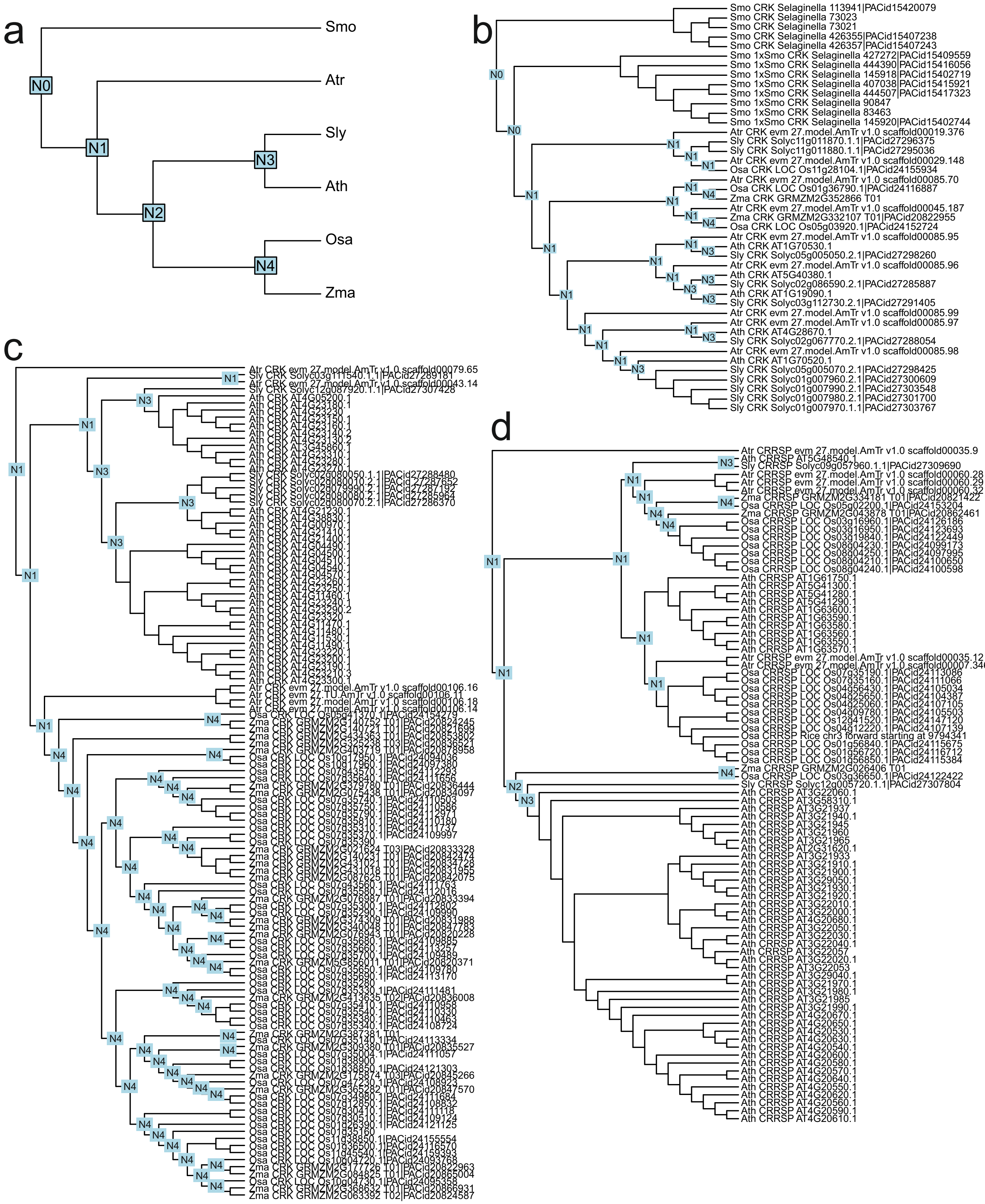


**Figure S14. Gene tree reconciliation of the five species used in segmental duplication analyses. a)** Species tree of the five species in the analyses. The phylogeny was downloaded from PhyloT, and the node labels indicate the speciation event ID. These IDs are used in Figures S15b-d). The reconciled gene trees were estimated with DLCpar for **b)** bCRKs, **c)** vCRKs and **d)** ddCRRSPs. The node labels provide the timing of the split by referring to the speciation event ID in the species tree (Figure S15a). *Selaginella moellendorffii* was used as outgroup.

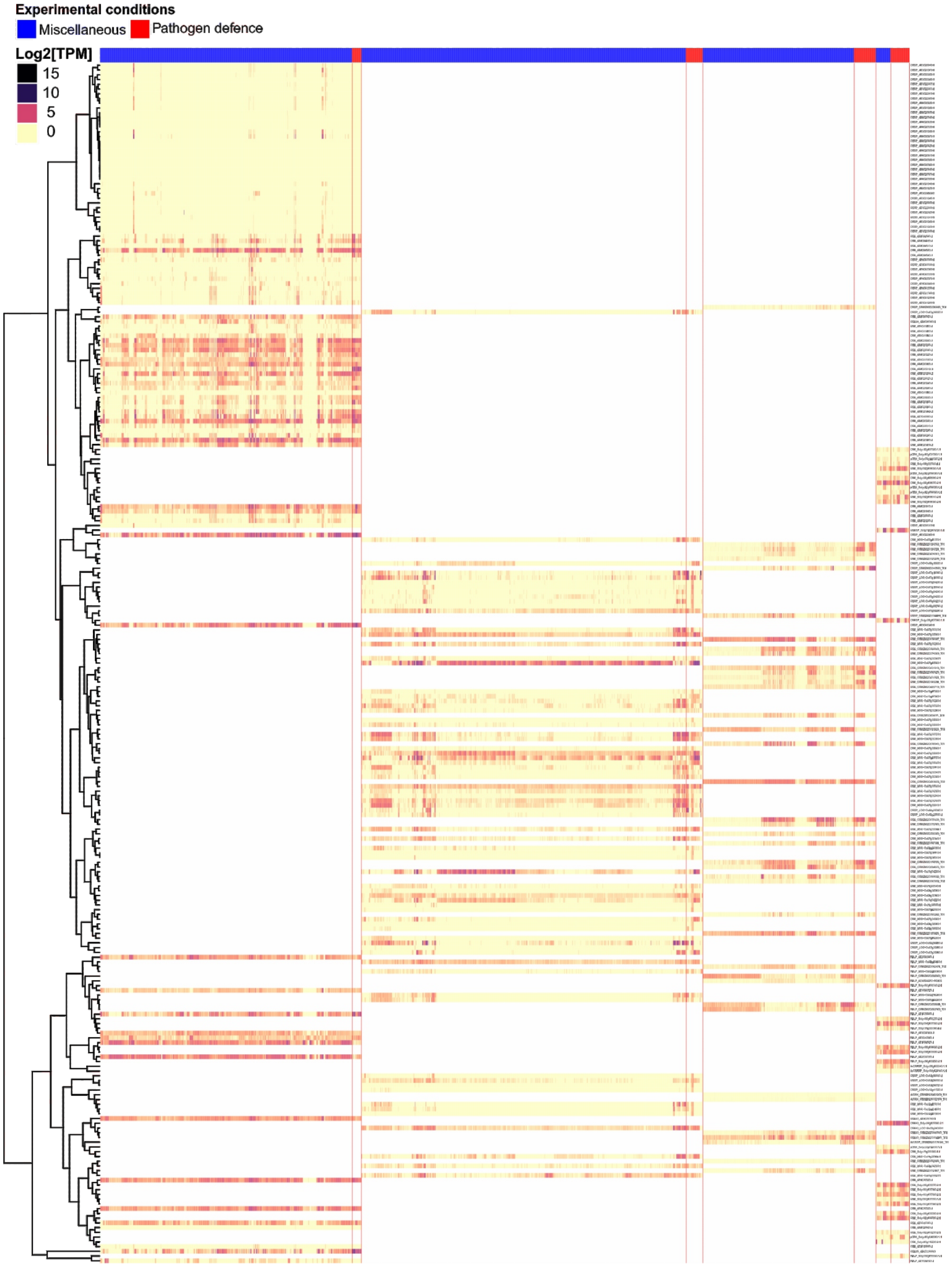


**Figure S15. The DUF26 genes show transcriptional response to several stress treatments.** Heatmap illustrating transcriptional response of DUF26 genes from *Arabidopsis thaliana*, *Oryza sativa*, *Zea mays* and *Solanum lycopersicum*. The dendrogram shows a phylogenetic tree of the 253 DUF26-containing genes (rows) in the four species. The columns represent the RNAseq experiments from Sequence Read Archive (see Table S4; accession numbers not shown here for clarity), categorized into pathogen defence (red highlight) and miscellaneous (blue). The heatmap colors represent the $\log_2$(TPM) values, as illustrated by the color key. The NA values are displayed with white color.
