## Supplementary material for "Mechanistic insights into the evolution of DUF26-containing proteins in land plants"

Table S2 **Data collection, phasing and refinement statistics**

| | **PDLP5** *sulfur SAD** | **PDLP5** *native** | **PDLP8** *native** |
|---|---|---|---|
| **Data collection** | | | |
| Space group | *P1* | *P1* | $P3_2\,2\,1$ |
| Cell dimensions | | | |
| *a*, *b*, *c* (Å) | 41.81, 48.05, 62.24 | 41.76, 47.97, 62.19 | 143.86, 143.86, 59.72 |
| α, β, γ (°) | 97.68, 102.74, 99.90 | 97.70, 102.72, 99.86 | 90, 90, 120 |
| Resolution (Å) | 48.91 – 2.3 (2.36 – 2.30) | 37.77 – 1.29 (1.32 – 1.29) | 19.91 – 1.95 (2.02 – 1.95) |
| $R_{meas}$# | 0.004 (0.10) | 0.049 (1.29) | 0.319 (3.62) |
| $I/\sigma I$# | 77.5 (33.5) | 12.5 (1.1) | 8.82 (1.0) |
| *CC (1/2)*# | 99.9 (99.9) | 99.9 (55.7) | 99.8 (64.7) |
| Completeness (%)# | 91.2 (83.6) | 94.5 (89.4) | 100.0 (99.9) |
| Redundancy# | 28.2 (28.5) | 3.6 (3.7) | 20.1 (20.1) |
| **Refinement** | | | |
| Resolution (Å) | | 37.77 – 1.29 | 19.91 – 1.95 |
| No. reflections | | 103,379 | 49,227 |
| $R_{work}/R_{free}$$ | | 0.18/0.21 | 0.23/0.26 |
| No. atoms | | | |
| protein | | 3,201 | 4,821 |
| ligands | | 153 | 28 |
| solvent | | 310 | 193 |
| Res. B-factors$ | | | |
| protein | | 23.1 | 42.7 |
| ligands | | 47.5 | 70.1 |
| solvent | | 32.3 | 43.6 |
| R.m.s deviations$ | | | |
| Bond lengths (Å) | | 0.012 | 0.015 |
| Bond angles (º) | | 1.58 | 1.36 |
| Ramachandran favored (%) | | 98.5 | 97.24 |
| Ramachandran allowed (%) | | 1.23 | 2.76 |
| Ramachandran outliers (%) | | 0.25 | 0 |
| PDB - ID | | **6GRE** | **6GRF** |

#as defined XDS[11] or $in Refmac5[17], respectively. *Data were collected from one crystal per experiment.
