## Supplementary material for "Mechanistic insights into the evolution of DUF26-containing proteins in land plants"

| Species | Genome annotation version |
|---|---|
| *Amborella trichopoda*[1] | v1.0 |
| *Aquilegia coerulea*[*] | v1.1 |
| *Arabidopsis lyrata*[2] | v1.0 |
| *Arabidopsis thaliana*[3,4] | TAIR10 |
| *Betula pendula*[5] | v1.0 |
| *Brachypodium distachyon*[6] | v3.0 |
| *Capsella rubella*[7] | v1.0 |
| *Chlamydomonas reinhardtii*[8] | v5.5 |
| *Coccomyxa subellipsoidea*9 | v2.0 |
| *Cucumis sativus** | v1.0 |
| *Hordeum vulgare*[10] | v2.2 |
| *Klebsormidium flaccidum*[11] | v1 |
| *Marchantia polymorpha*[12] | v3.1 |
| *Medicago truncatula*[13,14] | Mt4.0v1 |
| *Micromonas pusilla*[15] | v3.0 |
| *Nelumbo nucifera*[16] | v1 |
| *Oryza sativa*[17,18] | v7 |
| *Ostreococcus lucimarinus*[19] | v2.0 |
| *Physcomitrella patens*[20] | v3.0 |
| *Picea abies*[21] | v1.0 |
| *Populus trichocarpa*[22] | v3.0 |
| *Prunus persica*[23] | v1.0 |
| *Selaginella moellendorffi*[i24] | v1.0 |
| *Solanum lycopersicum*[25] | iTAG2.4 |
| *Solanum melongena*[26] | r2.5.1 |
| *Solanum tuberosum*[27] | v3.4 |
| *Sorghum bicolor*[28] | v2.1 |
| *Spirodela polyrhiza*[29] | v1 |
| *Theobroma cacao*[30] | v1.1 |
| *Vitis vinifera*[31] | Genoscope.12X |
| *Volvox carteri*[32] | v2.0 |
| *Zea mays*[33] | AGPv3 |
